# OPENPichia: building a free-to-operate *Komagataella phaffii* protein expression toolkit

**DOI:** 10.1101/2022.12.13.519130

**Authors:** Dries Van Herpe, Robin Vanluchene, Kristof Vandewalle, Sandrine Vanmarcke, Elise Wyseure, Berre Van Moer, Hannah Eeckhaut, Daria Fijalkowska, Hendrik Grootaert, Chiara Lonigro, Leander Meuris, Gitte Michielsen, Justine Naessens, Charlotte Roels, Loes van Schie, Riet De Rycke, Michiel De Bruyne, Peter Borghgraef, Katrien Claes, Nico Callewaert

## Abstract

In the standard toolkit for recombinant protein expression, the yeast known in biotechnology as *Pichia pastoris* (formally: *Komagataella phaffii*) takes up the position between *E. coli* and HEK293 or CHO mammalian cells, and is used by thousands of laboratories both in academia and industry. The organism is eukaryotic yet microbial, and grows to extremely high cell densities while secreting proteins into its fully defined growth medium, using very well established strong inducible or constitutive promoters. Many products made in *Pichia* are in the clinic and in industrial markets. *Pichia* is also a favoured host for the rapidly emerging area of ‘precision fermentation’ for the manufacturing of food proteins. However, the earliest steps in the development of the industrial strain (NRRL Y-11430/CBS 7435) that is used throughout the world were performed prior to 1985 in industry (Phillips Petroleum Company) and are not in the public domain. Moreover, despite the long expiry of associated patents, the patent deposit NRRL Y-11430/CBS 7435 that is the parent to all commonly used industrial strains, is not or no longer made freely available through the resp. culture collections. This situation is far from ideal for what is a major chassis for synthetic biology, as it generates concern that novel applications of the system are still encumbered by licensing requirements of the very basic strains. In the spirit of open science and freedom to operate for what is a key component of biotechnology, we set out to resolve this by using genome sequencing of type strains, reverse engineering where necessary, and comparative protein expression and strain characterisation studies. We find that the industrial strains derive from the *K. phaffii* type strain lineage deposited as 54-11.239 in the UC Davis Phaff Yeast Strain collection by Herman Phaff in 1954. This type strain has valid equivalent deposits that are replicated/derived from it in other yeast strain collections, incl. in ARS-NRRL NRRL YB-4290 (deposit also made by Herman Phaff) and NRRL Y-7556, CBS 2612 and NCYC 2543. We furthermore discovered that NRRL Y-11430 and its derivatives carry an ORF-truncating mutation in the *HOC1* cell wall synthesis gene, and that reverse engineering of a similar mutation in the NCYC 2543 type strain imparts the high transformability that is characteristic of the industrial strains. Uniquely, the NCYC 2543 type strain, which we propose to call ‘OPENPichia’ henceforth, is freely available from the NCYC culture collection, incl. resale and commercial production licenses at nominal annual licensing fees^1^. Furthermore, our not-for-profit research institute VIB has also acquired a resale/distribution license from NCYC, which we presently use to openly provide to end-users our genome-sequenced OPENPichia subclone strain and its derivatives, i.e., currently the highly transformable *hoc1*^*tr*^ and the *his4* auxotrophic mutants. To complement the OPENPichia platform, a fully synthetic modular gene expression vector building toolkit was developed, which is also openly distributed, for any purpose. We invite other researchers to contribute to our open science resource-building effort to establish a new unencumbered standard chassis for *Pichia* synthetic biology.

## Introduction

Presently, a wide variety of microbial hosts is available for the production of recombinant proteins. However, for general laboratory use as well as biopharmaceutical protein manufacturing, a strong consolidation has taken place over the past years. *E. coli* remains the most-frequently used prokaryotic system for proteins of prokaryotic origin and for relatively simple stably folding proteins of eukaryotic origin, especially those with no or few disulphide bonds. On the other end, production in HEK293 or CHO cells is used for production of proteins of higher eukaryotic origin, affording *i*.*a*. complex-type N-glycosylation. The methylotrophic yeast known in biotechnology as *Pichia pastoris* (and formally classified as *Komagataella phaffii*) takes up the intermediate-complexity position in most protein expression lab’s toolkit, as it combines the easy cultivation and fast growth of a micro-organism with the presence of a eukaryotic secretory system and the ensuing ability to perform complex post-translational modifications such as N-glycosylation and strong capacity for the formation and isomerization of disulphide bonds^2–4^. Hundreds of papers are published every year reporting on the use of *Pichia* for protein production (and increasingly also for engineered metabolite production, incl. in the area of sustainable chemical building block manufacturing from methanol and even CO ^5^). Quite often, *Pichia*-produced proteins are developed by academic and industrial scientists alike with an eventual applied/commercial use in mind, being it as a therapeutic compound, an industrial biocatalyst^6,7^ or more recently, as a food ingredient.

Historically, in 1954 Herman Phaff deposited a methylotrophic yeast strain from a black oak tree (*Quercus*) in the Yosemite region^8^. The isolate was stored in the culture collection of the University of California at Davis as UCD-FST K-239, with formally equivalent type strain deposits in other culture collections as NRRL YB-4290, NRRL Y-7556, CBS 2612 and NCYC 2543. At this time, these isolated strains of Phaff could not be distinguished from similar strains isolated in 1919 by A. Guilliermond, and Herman Phaff categorized both together as a new species: *Pichia pastoris* (the genus *Pichia* was established half a century before, in 1904, by E. Hanssen^9^) (Figure 1). In 1995, *Pichia pastoris* was re-classified into the new genus of *Komagataella*, named after the Japanese scientist Kazuo Komagata as a tribute to his contributions to yeast systematics, in particular the methanol-assimilating yeasts^10^. In 2005, the two distinctly evolved isolates from Phaff and Guilliermond were divided into two separate species and renamed *Komagataella phaffii* and *Komagataella pastoris* by C. Kurtzman^11^, based on the sequencing of 26S rDNA. Consequently, the strain of Phaff (UCD-FST K-239, NRRL YB-4290, NRRL Y-7556, CBS 2612, and NCYC 2543) was now considered the type strain of the species *Komagataella phaffii*, while the strain of Guillermond (CBS 704 or NRRL Y-1603) was regarded as the type strain of the species *Komagataella pastoris*.

**Figure 1.**
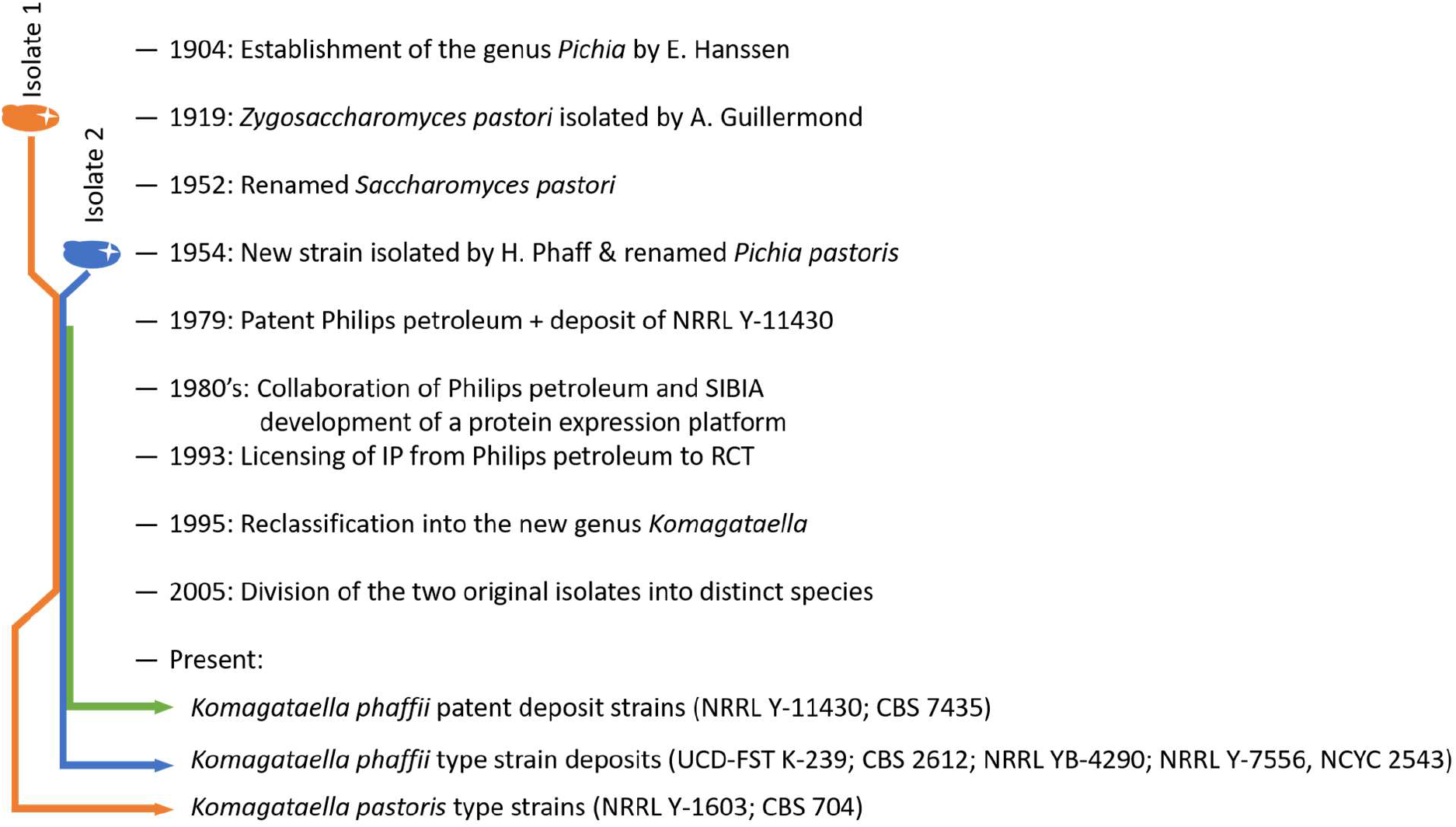
Schematic timeline of the history of the Komagataella species and the available type strain deposits and patent strain deposits.

By the 1970’s, these yeast species that can utilize methanol as the sole carbon source^12–14^ sparked the interest of Phillips Petroleum Company, since they had a vast supply of cheap methane gas (a by-product of their oil refinement process), which can be easily oxidized to methanol by chemical oxidation. Hence, they explored the available methylotrophic yeast species through procedures that are not in the public domain, and selected a *Pichia pastoris* to grow on the synthesized methanol and produce a single cell protein source for animal feed using fermentation. The application was patented in 1980 (with the earliest priority application on April 12^th^, 1979)^15^, which required the strain to be deposited once more in public culture collections and it became known as NRRL Y-11430 and CBS 7435. Which exact strain was deposited, is not described in the patent, nor available in the public domain. It was already assumed that it could be either the isolate from Guillermond (NRRL Y-1603) or that of H. Phaff (NRRL Y-4290) (GRN 737). However, recent efforts of genomic sequencing show that the genetic differences between both lineages are large enough to conclude that the latter was used^16,17^. In the early 80’s, Phillips Petroleum Company contracted with the Salk Institute Biotechnology/Industrial Associates (SIBIA) to develop the organism for recombinant protein production, based primarily on the extremely strong and tightly regulated Alcohol Oxidase 1 promoter (pAOX1). In this context, NRRL Y-11430 derived strains were generated with auxotrophies, such as the GS115 strain, which is a *his4* auxotrophic mutant obtained through nitrosoguanidine mutagenesis of NRRL-Y11430^18^, and the X33 strain that is a *HIS4* complemented strain deriving from GS115, likely via an intermediate^18–20^. In 1993, Phillips Petroleum sold its patent position in the *Pichia* system to Research Corporation Technologies (RCT, Tucson, AZ)^21^, apparently including the strain patent deposits associated to this, and their derived strains. Unfortunately, NRRL Y-11430 is not distributed anymore, likely because the patentee is no longer under an obligation to provide the strain, given that the associated patent(s) have expired almost 20 years ago. Indeed, the NRRL Y-11430 in the Agricultural Research Service culture collection (ARS-NRRL) is no longer available, and the same is the case for the CBS 7435 strain in the collection of the Westerdijk Fungal Biodiversity Institute (CBS). Also, it appears that the strain and its popular derivatives, incl. their distribution, are still controlled by the original patent holder. This occurs through the enforcement of Material Transfer Agreements (MTAs) that were associated to obtaining the strain from the NRRL culture collection during the time it was maintained as a patent deposit, or conditions of sale associated to obtaining derivative strains through the commercial distribution by Invitrogen (currently a brand of Thermo Fisher Scientific) as part of *Pichia* expression system kits. To our knowledge, the NRRL Y-11430 parental industrial strain can currently only be obtained at the American Type Culture Collection (ATCC 76273) under a similarly restrictive MTA. Because of the RCT-mandated distribution of *Pichia* expression technology kits by Invitrogen, for more than two decades, thousands of protein expression labs have extensive experience with the GS115 family of strains that were included in these kits^22,23^ and the currently approved biopharmaceuticals are manufactured by use of the NRRL Y-11430 industrialized strain lineage. Many researchers are not fully aware of the restrictions to strain distribution and potential applied use downstream of discoveries and inventions made with them. Without prejudicing the legal aspects of some of this, which is outside of our field of competence, as scientists we find it entirely unsatisfactory that the very basis of a critical mainstay of academic and industrial biotechnology remains not simply openly accessible exceedingly long after the original patents have expired. Given that *Pichia* takes such an ever more prominent place in the toolkit for diverse sectors of synthetic biology and biotechnology, we believe that it is long overdue for an open-access alternative strain platform to be set up by the community. To continue to take benefit of the extensive regulatory agency acquaintance with *Pichia*, it is important to achieve this by use of the same parental strain lineage as used in the strains that were industrialized so far.

To achieve this goal, we and others have recently turned to the *Komagataella phaffii* type strains that are present in culture collections throughout the world, armed with now highly affordable genome resequencing, to try and identify the original isolate from nature that the Phillips Petroleum researchers used in their derivation of NRRL Y-11430. In an exploration in the largest US yeast strain collection (ARS-NRRL), the Love lab at MIT found that two deposits indicated by the culture collection as equivalent *Komagataella phaffii* type strains (NRRL YB-4290, directly deposited by Herman Phaff, and NRRL Y-7556), are genetically identical. Furthermore, in comparison to this type strain, in their analysis they found only 1 point mutation leading to a truncation of the Rsf2p open reading frame in NRRL Y-11430^24^, establishing the type strain lineage of this industrial strain. In this study, the NRRL type strains were found to have problems with easily generating high copy number transformants and the authors recommended to keep using the industrial NRRL Y-11430 strain background for recombinant protein production, especially for expressions under methanol control. The NRRL YB-4290 strain is also present in the UC Davis Phaff Yeast Culture Collection as its deposit made by Herman Phaff in 1954, and numbered 54-11.239 (referred to as UCD-FST K-239 in the NRRL entries). It is documented in the collections as having been isolated at Mather, Central Sierra Nevada, California, USA, from the exudate flux of a black oak (*Quercus kelloggii*). Given its origin, no Access and Benefit Sharing restrictions apply to its use for any purpose (Nagoya Protocol or Convention on Biological Diversity). This type strain was also deposited in European culture collections: at the Westerdijk Fungal Biodiversity Institute (CBS, Delft, The Netherlands, deposit CBS 2612) and at the National Collection of Yeast Cultures (Norwich, UK, deposit NCYC 2543). In our present study, we started out by analysing the genome of the four type strains. We then focused on the NCYC 2543 strain, as the NCYC collection uniquely provides standard very affordable distribution and commercial use licenses for its strains and derivatives thereof, fulfilling our requirements for a community open-access *Pichia* platform strain. Different from all other culture collections where the *K. phaffii* type strain deposits are found, technology developers can acquire a distribution license to end-users for their newly generated materials, or have the choice of also depositing these in the NCYC collection for full open access. Hence, we propose to nickname this NCYC 2543 type strain deposit as ‘OPENPichia’. In an effort to establish confidence in the OPENPichia strain as a new standard open-access *Pichia* chassis for use by the research and biotechnological industry community, we set out on a comprehensive characterization of its genome, growth characteristics, transformation efficiency and recombinant protein expression, in comparison to the NRRL Y-11430 industrial strain. We discovered a key frameshift mutation in the *HOC1* gene of the industrial strain that enhances conduciveness to transformation, and introduced this mutation in OPENPichia using open-source genome engineering technology, to overcome this important limitation. Further characterization of this OPENPichia-*hoc1*^*tr*^ strain in comparison to NRRL Y-11430 in terms of growth rate, cell wall density and cell sensitivity to cell envelope-binding chemicals confirmed it to be indistinguishable phenotypically from the industrial strain. As set forth below, we complement this OPENPichia strain set with an OPENPichia modular protein expression vector toolkit, entirely made up of synthetic DNA to avoid any third-party MTAs, and following the Golden Gate cloning standard for compatibility with complementary toolkits from other *Pichia* developer labs^25^. The basic NCYC 2543 OPENPichia strain is available from NCYC, incl. straightforward and low-cost distribution licenses for labs that develop novel strains. Our own derived OPENPichia strains described in this paper are openly available for end-users (incl. for royalty-free *Pichia*-made commercial product manufacturing) under such distribution license that our not-for-profit research institution VIB obtained from NCYC. OPENPichia vector cloning materials are openly distributed for any utilization purpose in association with the public plasmid collection of the Belgian Coordinated Collection of Microorganisms.

## Results

### The genomes of the *K. phaffii* type strains and commercial strains are virtually identical

To evaluate alternative *K. phaffii* strains at the genomic level, we deeply (avg. 180x genome coverage) resequenced the genome of several type strains available at culture collections (i.e., NRRL YB-4290 / NRRL Y-7556 / CBS 2612 / NCYC 2543) in comparison to the NRRL Y-11430 industrial strain (**Error! Reference source not found**.) (Supplementary Table 1 and 2). The reads were mapped against the published reference genome of the CBS 7435 strain incl. the mitochondrial genome sequence and that of the two *K. phaffii* linear killer plasmids^26^. CBS 7435 is a deposit of the NRRL Y-11430 industrial strain in the CBS culture collection (Delft, The Netherlands). To this end, the Breseq^27^ software package was used in consensus mode. There was some variation in overall GC content, which can be attributed to the varying amount of mitochondrial DNA and the presence of the two killer plasmids in some of the strains, as these have an average GC content of 24%, 28% and 29%, respectively (unlike the nuclear chromosomes, which have an average GC content of 41%). Indeed, the read alignment showed that the proportion of reads that mapped to the mitochondrial genome varied between 10 and 47%.

**Table 1.** Strains used in this publication. * Type strain; Abbreviations: UCD: University of California, Davis, USA; CBS: Centraalbureau voor Schimmelculturen, currently known as Westerdijk Fungal Biodiversity Institute, The Netherlands; NCYC: National Collection of Yeast Cultures, UK; NRRL: Northern Regional Research Laboratory, currently known as Agricultural Research Service (ARS), USA; NTG: nitrosoguanidine.

| Strain ID | Origin of strain isolate | Source | Original depositor |
| --- | --- | --- | --- |
| CBS 2612 * | *Quercus kelloggii* (California, USA) | CBS culture collection | HJ Phaff, UCD |
| NCYC 2543 * | *Quercus kelloggii* (California, USA) | NCYC culture collection | CBS |
| NRRL YB-4290 * | *Quercus kelloggii* (California, USA) | NRRL culture collection | HJ Phaff, UCD |
| NRRL Y-7556 * | *Quercus kelloggii* (California, USA) | NRRL culture collection | D. Yarrow, CBS |
| UCD FST K-239 * | *Quercus kelloggii* (California, USA) | UCD culture collection | HJ Phaff, UCD |
| NRRL Y-11430 | Most likely a subclone of the type strain | NRRL culture collection | Patent deposit<br>Phillips Petroleum Company (USA) |
| GS115 | NTG-mutagenized derivative of NRRL Y-11430 | Internal VIB culture collection |  |
| NCYC 2543 *his4* | Mutagenesis | This publication |  |
| NCYC 2543 *hoc1<sup>tr-1</sup>* | Mutagenesis | This publication |  |
| NCYC 2543 *hoc1<sup>tr-2</sup>* | Mutagenesis | This publication |  |

The proportion of reads originating from the two killer plasmids, varied between 0% and 9% (Supplementary Table 2). The *K. phaffii* killer plasmids are linear autonomously replicating DNA fragments of 9.5 kb and 13.1 kb^19^ that encode exotoxins that can kill other yeast cells^17,19^. These toxins may also be toxic to cells from the same species, in case these have lost resistance to the toxin, and this phenomenon is therefore undesired in large scale culturing of yeast as it may reduce the overall viability of the culture. To verify the presence of such killer plasmids in the tested strains, their sequences, as reported by Sturmberger *et. al*., were included in the reference for the alignment of the reads^26^. Surprisingly, the killer plasmids could not be detected in both the CBS 2612 and NCYC 2543 strain, while such reads were abundant for the analysed NRRL strains (NRRL YB-4290, Y-7556 and Y-11430) (Table 2). According to the strain history, the NRRL YB-4290 and CBS 2612 strains were deposited by Phaff, while the NRRL Y-7556 strain was a redeposit of CBS 2612 by D. Yarrow (CBS). Since the daughter strain (NRRL Y-7556), still has these killer plasmids, the mother strain (CBS 2612) should have had them as well, leading to the conclusion that these linear plasmids are easily lost while propagating *K. phaffii* strains. In our lab as well, several NRRL Y-11430 descendants were identified that have lost these killer plasmids upon the standard microbial practice of single colony streaking (unpublished data). Also, the commercial strain GS115, does not have any killer plasmids^18^.

**Table 2.** Overview of the functional mutations of the analysed K. phaffii strains compared to the CBS 7435 (eq. strain deposit of NRRL-Y-11430) reference genome, non-functional SNPs and indels are between brackets. Functional mutations are those resulting in a SNP or indel in a gene.

| Strain Designation | SNP | Indel | Total | Mutations/Mbp | Killer plasmids copy number (KP1/KP2) |
| --- | --- | --- | --- | --- | --- |
| NRRL Y-11430 | 0 (2) | 2 (18) | 2 (20) | 2.3 | 21/15 |
| NRRL YB-4290 | 2 (2) | 3 (20) | 5 (22) | 2.9 | 140/82 |
| NCYC 2543 | 3 (5) | 3 (18) | 6 (23) | 3.1 | none detected |
| CBS 2612 | 3 (4) | 3 (19) | 6 (23) | 3.1 | none detected |
| NRRL Y-7556 | 3 (3) | 3 (21) | 6 (24) | 3.2 | 138/86 |

Using a Maximum Likelihood method and the Hasegawa-Kishino-Yano model, a phylogenetic tree based on the nuclear genome was computed to visualize the genetic distances between the sequenced strains and a variety of other *K. phaffii* strains whose genomes were published previously^18,24,28^. All *K. phaffii* type strains are strongly clustered together with the NRRL Y-11430 and CBS 7435 strains, as well as other close relatives (Figure 2). Thus, these data support previous literature^24,29^, where it was hypothesized that all these strains are derived from the same isolate, originally isolated by Phaff^28^.

**Figure 1:**
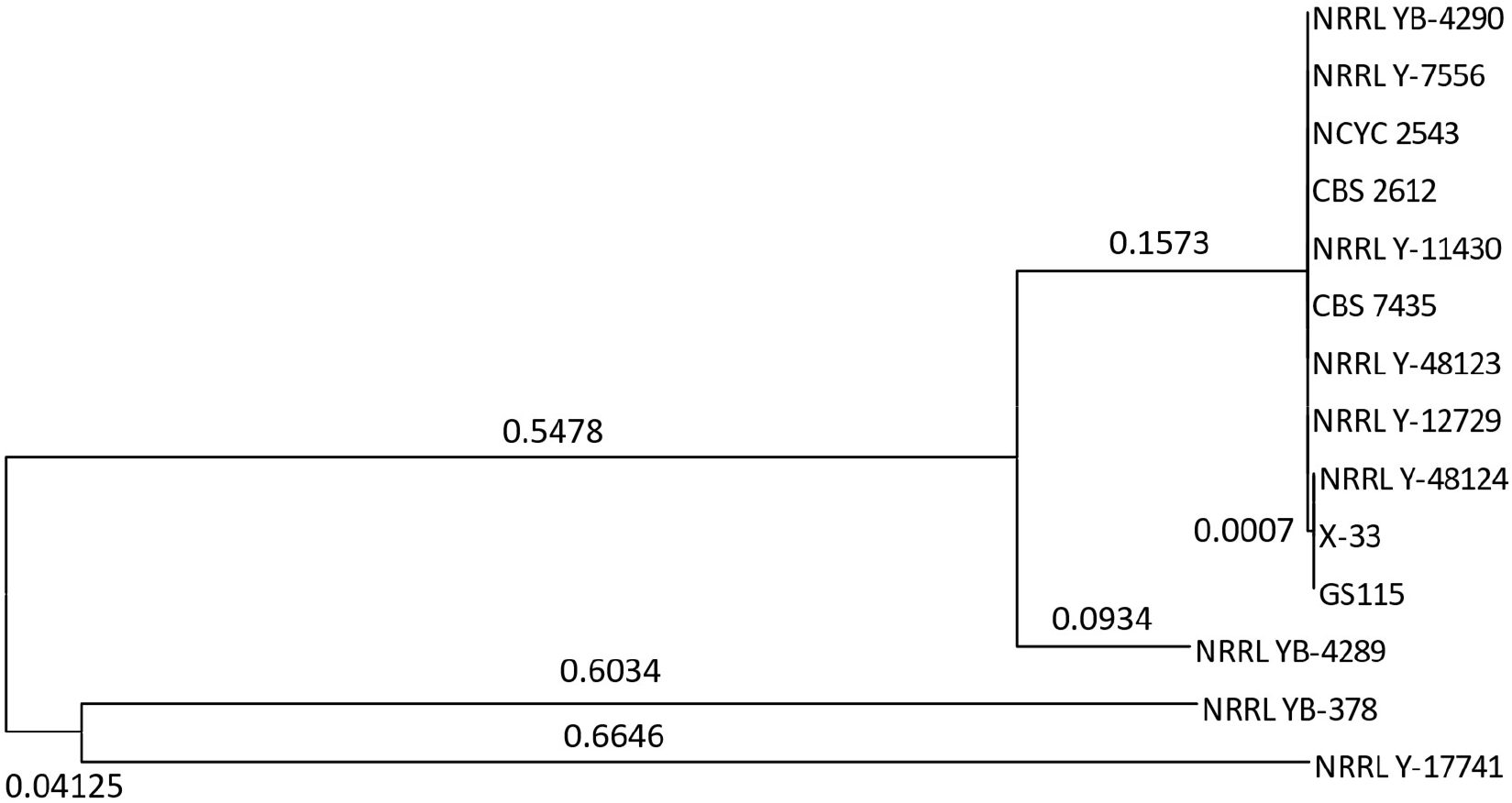
Phylogenetic tree of the K. phaffii strains with node lengths. The tree was constructed using a Maximum Likelihood method and Hasegawa-Kishino-Yano model. Node lengths of less than 0.0001 were neglected.

Based on the resequencing data, we identified single nucleotide polymorphisms (SNPs) and short insertion-deletions (indels) (Table 2). First comparing the mutation calling of strain NRRL Y-11430 with the reference genome of the equivalent-deposit CBS 7435 strain reference revealed only one protein-coding difference in one hypothetical protein and about 20 intergenic/intronic/silent exonic differences. All these alterations were also identified in the type strains, indicating that these are likely to be considered as the wild type *K. phaffii* genotype and that the CBS 7435 reference genome was most likely miscalled at these sites. The same holds for two insertions that were observed in each of the sequenced strains in a region annotated to encode for a hypothetical papain-like cysteine protease. This region appears to have been difficult to sequence in previous experiments, as also in the datasets associated with a previous type strain genome sequencing study^24^, we found only a few reads mapping to this area, which were accompanied by many mutations, low quality bases (Phred scores of <28) and low overall mapping quality score (<20). Note that the type strain deposits of the different culture collections also differ from one another each at one other coding sequence-altering genomic position and a few non-coding ones, likely reflecting drift due to background mutational rate during strain propagation (Table 2 and 3). We then focused on the very few protein-coding alterations that consistently distinguish the industrial strain NRRL Y-11430 from the equivalent type strain deposits NRRL YB-4290, NRRL Y-7556, CBS 2612, and NCYC 2543. Three coding sequence altering mutations (in *SEF1, RSF2*, and *HOC1*) were shared by all type strains but were absent in the NRRL Y-11430 strain. Our re-analysis of the raw sequencing reads from a previous genome analysis of deposits NRRL YB-4290 and NRRL Y-7556 confirmed the presence of these mutations and their absence in the NRRL Y-11430 strain also based on these datasets^24^. Data quality in the area of the *SEF1* and *HOC1* mutation is rather poor in these earlier datasets, and the authors only detected the mutation in *RSF2*^24^. Since all three mutations are shared by the type strains, it must be concluded that they represent the original state of *K. phaffii*, and that NRRL Y-11430 is actually the mutant strain at these three loci.

**Table 3.**
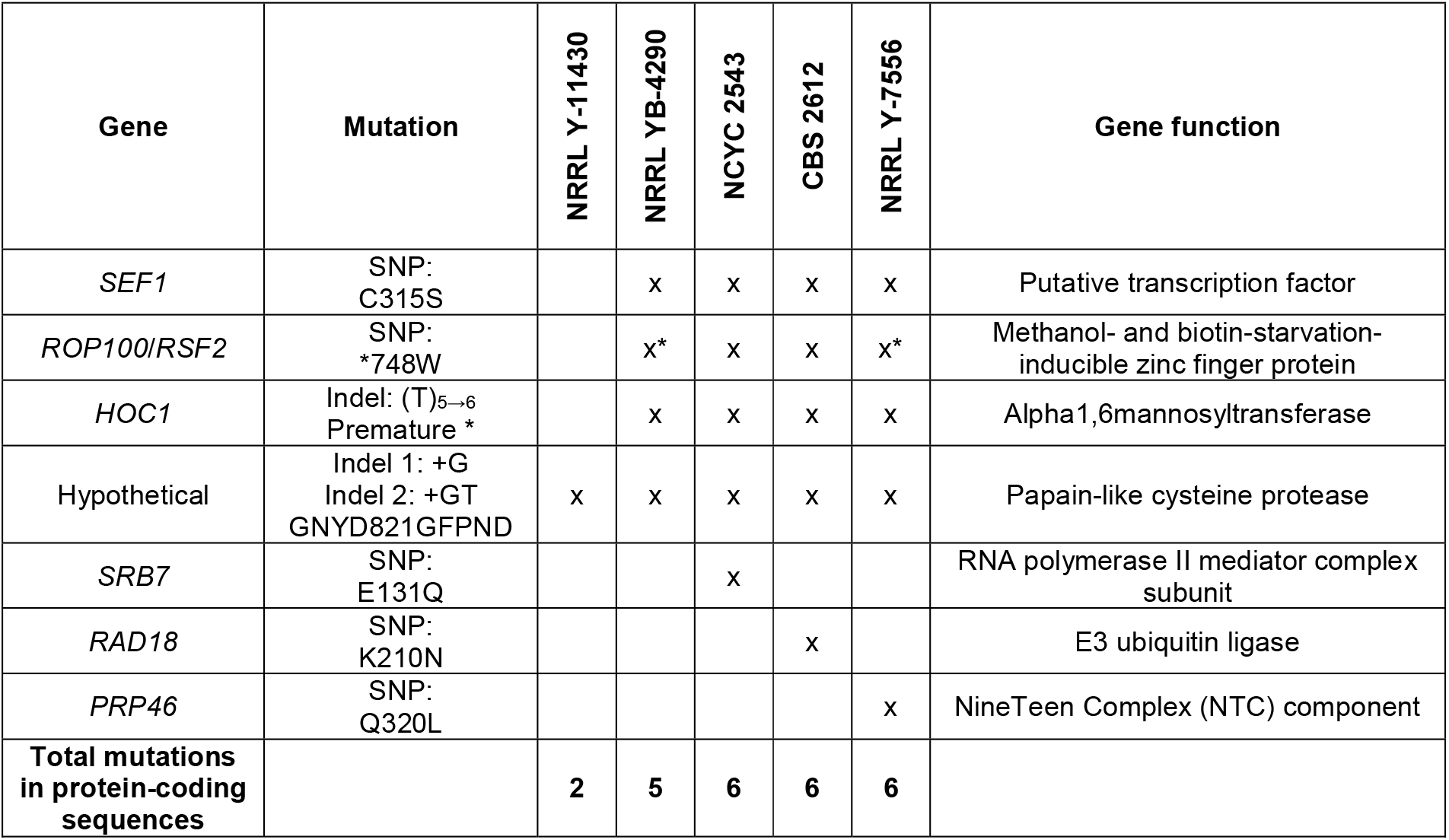
Summary of the coding mutations in the Philips Petroleum strain and the type strains, compared to the CBS 7435 reference genome. The exact mutations are mentioned in the second column, with the CBS 7435 amino acid as reference amino acid; although the original genetic makeup is as present in the type strains, and it is the CBS 7435/NRRL Y-11430 that mutated. The mutations indicated with an asterisk were also reported by Brady et. al.^24^. All mutations found in these strains are concentrated in eight locations. The mutations in SEF1, ROP100/RSF2, and HOC1 are shared by all type strains. Two mutations in a gene encoding for a hypothetical protein were found in all sequenced genomes. The other mutations (in SRB7, RAD18 and PRP46) are present in one of the type strains, NCYC 2543, CBS 2612 and NRRL Y-7556, respectively.

### Three mutations in protein-coding regions in the NRRL Y-11430/CBS 7435 industrial strain lineage vs. the type strain deposits

*SEF1* encodes for a putative transcription factor (UniProt ID F2QV09) and the observed SNP causes a S315C mutation in NRRL Y-11430 as compared to the type strains. *RSF2* encodes for a transcription factor which is involved in methanol- and biotin-starvation (UniProt ID F2QW29). The observed mutation is a SNP which causes the introduction of a stop codon (W748*) in NRRL Y-11430, resulting in the deletion of 183 amino acids from the C-terminus of the protein. The non-truncated Rsf2p which is found in the *K. phaffii* type strains, is very similar to its homolog in *S. cerevisiae*^30^, additionally indicating that this was the original genomic state, as previously reported^24^. *HOC1* (<u>h</u>omolog of *<u>OC</u>H<u>1</u>*) encodes for an α-1,6-mannosyltransferase (UniProt ID F2QVW2) involved in the synthesis of cell wall mannan and is part of the M-PolII complex^31^. Here, the industrial strain NRRL Y-11430 has a single base pair deletion in a poly-A stretch (at bp 755 of the 1191 bp long coding sequence), as compared to the type strains. This causes a frameshift and premature stop codon, resulting in a C-terminally truncated protein (274 versus 398 amino acids), with the last 22 amino acids up to the new stop codon being different from the type strain sequence. The indel in the homopolymer was confirmed by Sanger sequencing (data not shown). In parallel to our work, the same mutation was identified by the lab of Kenneth Wolfe (UC Dublin), as the phenotype-causative mutation for a quantitative trait locus (QTL) that yielded 2-3 fold higher secretion of a reported beta-glucosidase (personal communication, publication in press^29^). With this knowledge on which mutations could be involved in phenotypic differences between the type strains and the industrial strains, we set out for a comparison of characteristics important to the use of *Pichia* for recombinant protein production. Given that the NCYC 2543 deposit of the type strain was the only one for which the resp. culture collection openly provides both commercial and re-sale/strain distribution licenses as part of its standard business practice^1^, we further mainly focused on this deposit’s characteristics, and we called it OPENPichia.

### Strain comparison for growth rate, protein production and transformation efficiency

NCYC 2543/OPENPichia was compared to NRRL Y-11430 and GS115, both in terms of growth rate and their capacity for expressing recombinant proteins. GS115 is a *his4* auxotrophic mutant derived from NRRL Y-11430 by nitrosoguanidine mutagenesis and, depending on the analysis, its genome contains about 69^32^, 74^24^ or 71 (our unpublished data) mutations vs. that of its parent. To be able to evaluated the impact of *his4*-mediated histidine auxotrophy on strain characteristics, we generated an OPENPichia *his4* strain using the split-marker method^33,34^. No significant difference in growth rate between the type strains and NRRL Y-11430 could be observed (**Error! Reference source not found**.), but the GS115 strain grew significantly more slowly (one-way ANOVA, p=0.0034, Tukey test), as reported^24^. Interestingly, our histidine auxotrophic OPENPichia does not show the slower growth phenotype, demonstrating that this GS115 phenotype is not due to histidine auxotrophy, but rather must be due to one or more of the other nitrosoguanidine-induced mutations in its genome. As *his4* complementation is an often-used antibiotic-free selection marker, this unaffected growth rate in the OPENPichia *his4* strain is a useful feature.

To compare the protein production capacity of the strains, a selection of model proteins was produced in NRRL Y-11430 and OPENPichia (Table). Four proteins of very different types in use in biotechnology were chosen: a cytokine (GM-CSF), a redox enzyme (GaOx), a VHH-hFcα fusion (Cdiff-VHH-IgA), and a VHH-hFcγ fusion (CovidVHH-IgG). To enable recombinant expression, the two most commonly used off-patent *K. phaffii* promoters were tested: pGAP for strong constitutive and pAOX1 for strong methanol-inducible expression, respectively. Protein expression in *K. phaffii* is prone to clonal variations that can interfere with the comparison of expression capabilities between strains. Most of the variation is due to the integration site and the copy number of the construct^35^. To this end, a single-copy was targeted to the respective promoter regions in the genome, the copy number and integration site were confirmed by qPCR and integration-site specific PCR, and two independent clones of each setup were grown in triplicate.

For both the pGAP-driven and pAOX1-driven expressions, there is in general no major difference in protein production titres between the two hosts. However, there is a clear difference in cell density at harvest of the pGAP cultures, with OPENPichia growing to higher densities than NRRL Y-11430 (Supplementary Figure 1), whereas this is not the case on methanol. Additionally, in all cases, NRRL Y-11430 shows slightly more host cell proteins (HCPs) in the medium of the pGAP expressions (on limiting glucose), as visible on the SDS-PAGE gels (Figure 4), but not when grown on methanol. Both observations point towards a minimally increased cell lysis of the NRRL Y-11430 strain vs. the OPENPichia strain when grown on glucose.

**Figure 3.**
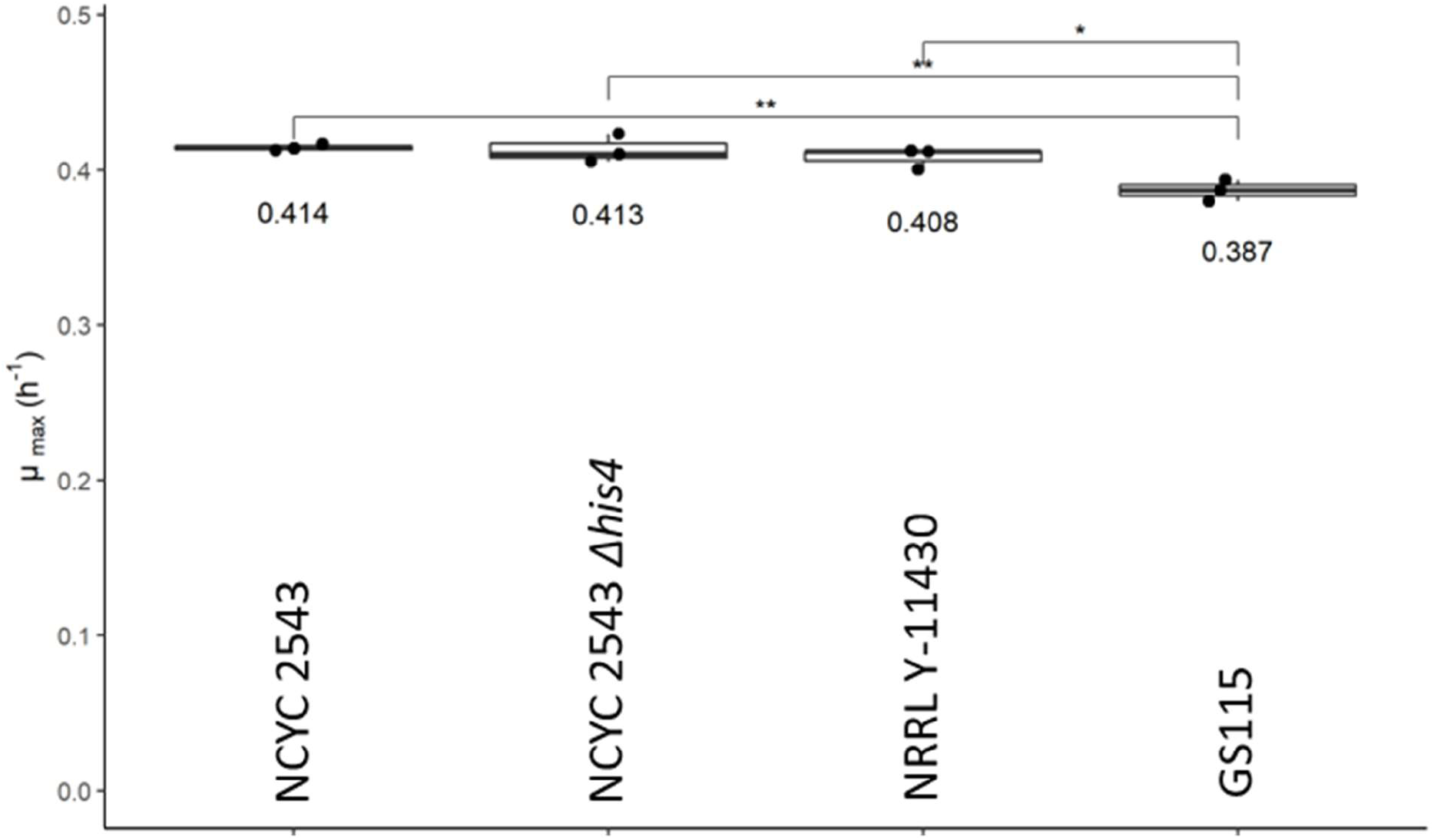
Maximum growth rate of the different Pichia strains, plotted individually as dots, as well as a boxplot with median value. A one-way ANOVA test showed significant influence of the strain on the growth rate (p = 0.0034). A post-hoc Tukey test shows that GS115 grows significantly slower than the other strains. The asterisks indicate the p-value of the Tukey test: **, 0.01>p>0.001; *, 0.05>p>0.01.

**Figure 4.**
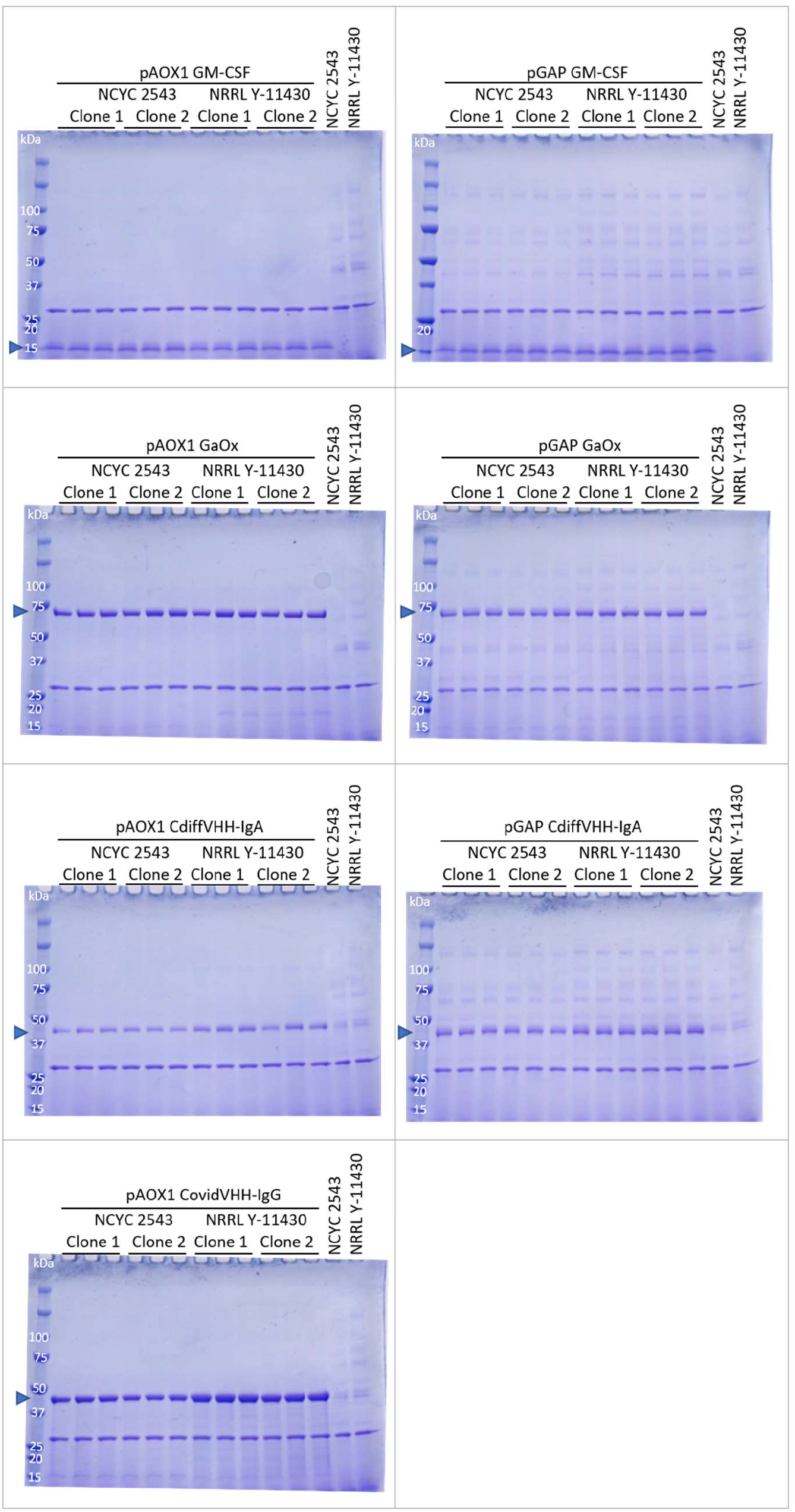
Expression comparison between NCYC 2543 and NRRL Y-11430. The proteins were expressed using the GAP or AOX1 promoter. As controls, both wild type strains were grown and analysed as well. Supernatant samples were treated with EndoH to remove N-glycans and samples were analysed on SDS-PAGE. EndoH is also visible on the gels at around 30 kDa.

### *HOC1*-truncation restores the transformation efficiency in NCYC 2543

Upon generating expression clones, a strongly reduced transformation efficiency was observed for OPENPichia as compared to NRRL Y-11430 (Figure 5C), as was also observed by others^24^. Out of the three consistent protein-coding differences between the type strains and NRRL Y-11430, we reasoned that the one in the *HOC1* open reading frame could be causative for the low transformation efficiency of the type strains, as the *S. cerevisiae* Hoc1p ortholog is an α-1,6-mannosyltransferase that is part of the mannan polymerase II complex in the yeast Golgi apparatus. The hypermannosyl-N-glycans, of which the backbone is synthesized by this mannan polymerase complex, form the outermost layer of the ascomycete cell wall, and carry most of their charge under the form of mannosylphosphate modifications on the side chains. *HOC1* deletion in *S. cerevisiae* is viable though the phenotyping and genetic interaction data available in the *Saccharomyces* Genome Database clearly indicate a cell wall stress response. As negatively charged plasmid DNA during transformation has to traverse the cell wall, a cell wall that presents less of a diffusional/charge barrier due to less mannan/mannosylphosphate density could explain the more highly transformable phenotype of NRRL Y-11430. Using the split-marker method, we introduced this single base pair deletion in the genome of OPENPichia, hence reverse engineering the genetic makeup of the NRRL Y-11430 strain (Figure 5A) in this locus. Two mutant versions were generated: NCYC 2543 *hoc1*^*tr*^-1, where only the single base pair was deleted, resulting in the truncated ORF and a Lox72-scar downstream of the novel stop codon; and NCYC 2543 *hoc1*^*tr*^-2, where additionally 115 bp downstream of the novel stop codon were removed (Figure 5B). The transformation efficiency was compared between the two wild type strains and the *hoc1*^*tr*^ mutants, using a plasmid encoding for a VHH under control of the *GAP* or *AOX1* promoter. The *HOC1*-truncation drastically increased the transformation efficiency of the OPENPichia strain, and even showed a 2-3-fold improvement as compared to the NRRL Y-11430 in this experiment (Figure 5C). In general, it was also observed that the transformation efficiency for pGAP-based plasmids is lower as compared to similar plasmids that are pAOX1-based, which we speculate is due to the metabolic burden imposed by constitutive GBP VHH production during recovery of the cells after transformation.

**Figure 5.**
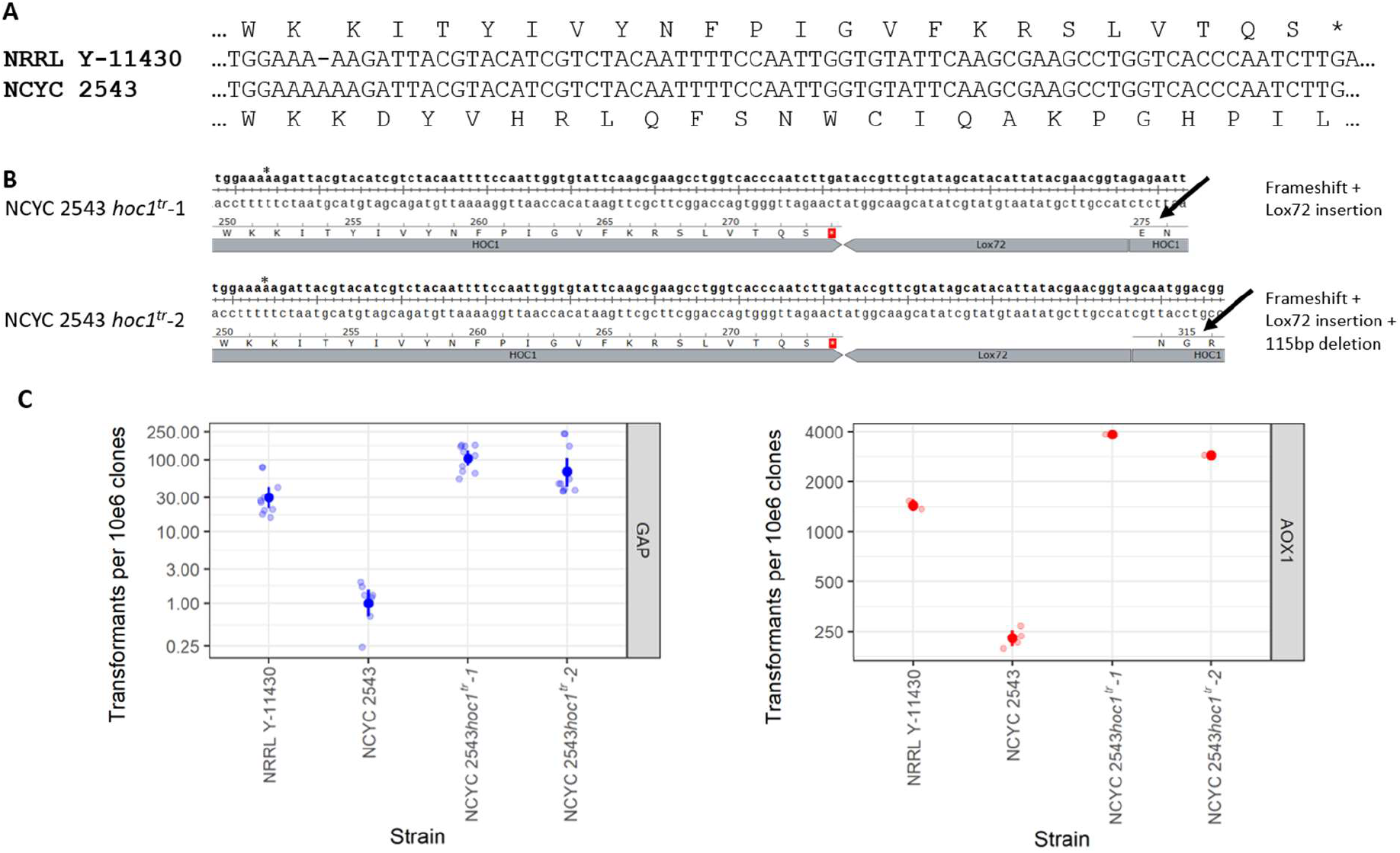
Overview of the HOC1 genome engineering strategy and the effect on transformation efficiency of the resulting strains. A. Alignment of a part of the HOC1 gene as present in NRRL Y-11430 vs. NCYC 2543, showing the frameshift resulting in a premature stop codon in the NRRL Y-11430. B. Resulting genomic HOC1 sequence upon split-marker-based gene editing. Two strategies were followed, where either the single base pair deletion (indicated with *) resulting in the Hoc1p truncation and a Lox72 scar is introduced downstream of the stop codon; or where an additional 115 bp deletion downstream of the resulting stop codon and Lox72 scar is introduced. C. Transformation efficiency in the two wild type strains and two HOC1-engineered strains, either using a pGAP-based plasmid (left) or a pAOX1-based plasmid (right). The analysis was performed as described in the Materials & Methods section.

### Characterization of the NRRL Y-11430, NCYC 2543 and NCYC 2543 *hoc1*^*tr*^ cell wall

These strains were further characterized in terms of their cell wall composition. First, the cell wall mannoprotein N-glycans were profiled by capillary electrophoresis^36^ after the cells were grown on glucose or glycerol (Supplementary Figure 2). This method mainly detects the lower-degree of polymerization N-glycans, and the profiles were very similar for all four strains, indicating that the pathway of synthesis of the mannan core was intact in all strains. This capillary electrophoresis method is however unsuited to the detailed profiling of the higher-polymerized mannan N-glycans, as there are so many isomeric structures formed that are all in part or completely resolved, such that they form one long trail of overlapping low-abundance peaks that is impossible to interpret. Most of the mannosylphosphate negative charge-imparting moieties are added to the mannan side branches of these long chains. They bind to cationic dyes such as Alcian Blue and hence the staining density of yeast cells with such dye reflects the density of these negative charges on the outermost layer of the cell wall. Indeed, as compared to the type strain NCYC 2543/OPENPichia, both NRRL Y-11430 and the two OPENPichia *hoc1*^*tr*^ mutants showed a reduced Alcian Blue staining intensity (Figure 6A). This is consistent with the reduced Alcian Blue staining of *S. cerevisiae hoc1* mutants^37,38^.

**Figure 6.**
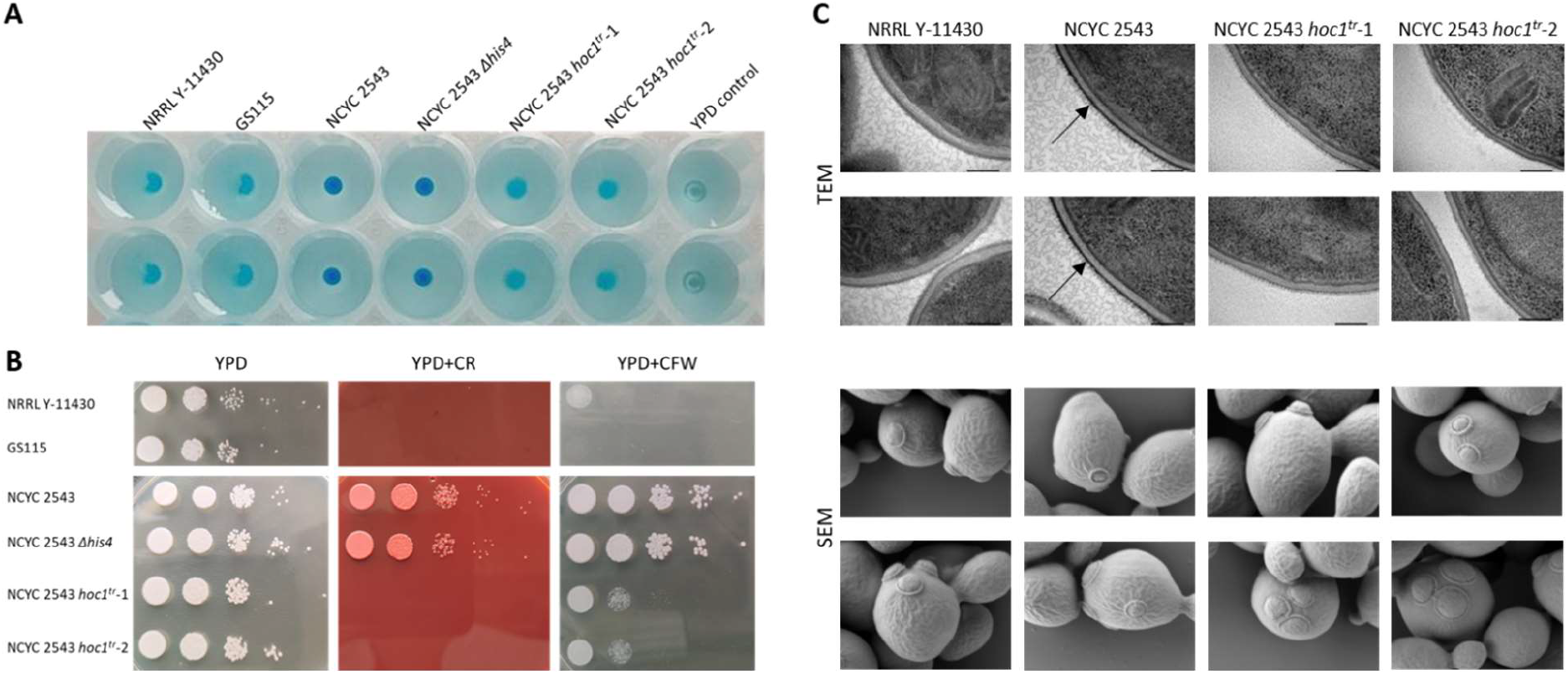
Characterization of the cell walls of NRRL Y-11430, NCYC 2543 and the two NCYC 2543 hoc1^tr^ mutants. A. Alcian blue staining of the strains to determine the density of negative charges at the yeast cell wall. Alcian blue is a cationic dye that binds negative charges at the cell wall. The more intense the blue staining of the cells, the more negative charge, i.e., mannosylphosphate moieties, are present on the glycan trees of the cell wall mannoproteins. Duplicate wells are shown per strain (vertical). B. Sensitivity of the strains towards Congo red and Calcofluor white, compared to growth on YPD agar, as an indicator for cell wall integrity. The plates were incubated for 3 days at 30 °C. C. Transmission electron microscopy (TEM) and scanning electron microscopy (SEM) images of the four strains. The increased electron density of the outermost layer, i.e., the cell wall is indicated with an arrow in the TEM images of the NCYC 2543 strain. Only two individual images per strain are shown.

To further investigate the cell wall integrity, resistance of the strains towards Congo red and Calcofluor white was analysed (Figure 6B) using previously reported methods^24,29^. The OPENPichia type strain is much more resistant than NRRL Y-11430 towards both dyes, a difference which is lost in the *HOC1*-truncated OPENPichia mutants. This indicates the importance of Hoc1p in cell wall integrity. We also performed transmission electron microscopy (TEM) using a freeze substitution technique that is optimized to pull in the osmium tetroxide membrane-staining contrast reagent as well as fixatives through the cell wall during the slow dehydration of the cells, which is then reversed in subsequent sample preparation. We observe a much stronger electron scattering at the outermost cell wall layer of the wild type OPENPichia type strain than for the other, *HOC1*-truncated strains (NRRL Y-11430 and OPENPichia *hoc1*^*tr*^) (Figure 6C). We interpret that this is caused by OsO4-accumulation at the mannan layer of the cell wall due to a stronger barrier to diffusion of the reagent in the wild type strain during freeze substitution. Scanning electron microscopy looked very similar for the four strains (Figure 6C), indicating no gross malformations. In summary, the data are consistent with the *hoc1*^*tr*^ mutation resulting in a relatively mild cell wall integrity deficiency, resulting in increased passage of plasmid DNA during transformation and, depending on the target protein, in some cases also increased production or cell wall passage of secreted recombinant proteins^29^.

### Strain comparison for growth rate and protein production of OPENPichia *hoc1*^*tr*^

The growth rate, as well as protein production capacity of the newly generated OPENPichia *HOC1*-truncated strains were compared to those of NRRL Y-11430 and wild type OPENPichia. No significant difference in growth rate is observed between the 4 tested strains (Figure 7A). As we earlier found most phenotypic difference between NRRL Y-11430 and wild type OPENPichia in terms of growth and HCP levels when the strains were grown on glucose, we tested pGAP-based protein expression for the different strains (Table 4), using an anti-GFP VHH (GBP) as test protein. To this end, 24 clones of each strain were screened, except for wild type OPENPichia, where only 12 transformants were obtained (Figure 7B and C). For pGAP-based GBP expression, the NCYC 2543 *hoc1*^*tr*^ strains outperform the NCYC 2543 strain, and NCYC 2543 *hoc1*^*tr*^-1 even outperforms NRRL Y-11430, although the differences are small and clonal distributions overlap. The result is in line with the observations made by the Wolfe lab (personal communication, publication in press^29^), where they observed that the truncated *HOC1* genotype in NRRL Y-11430 resulted in doubling of the secretion level of a beta-glucosidase under control of the pGAP promoter, vs. rather divergent *K. phaffii* NRRL strains.

**Figure 7.**
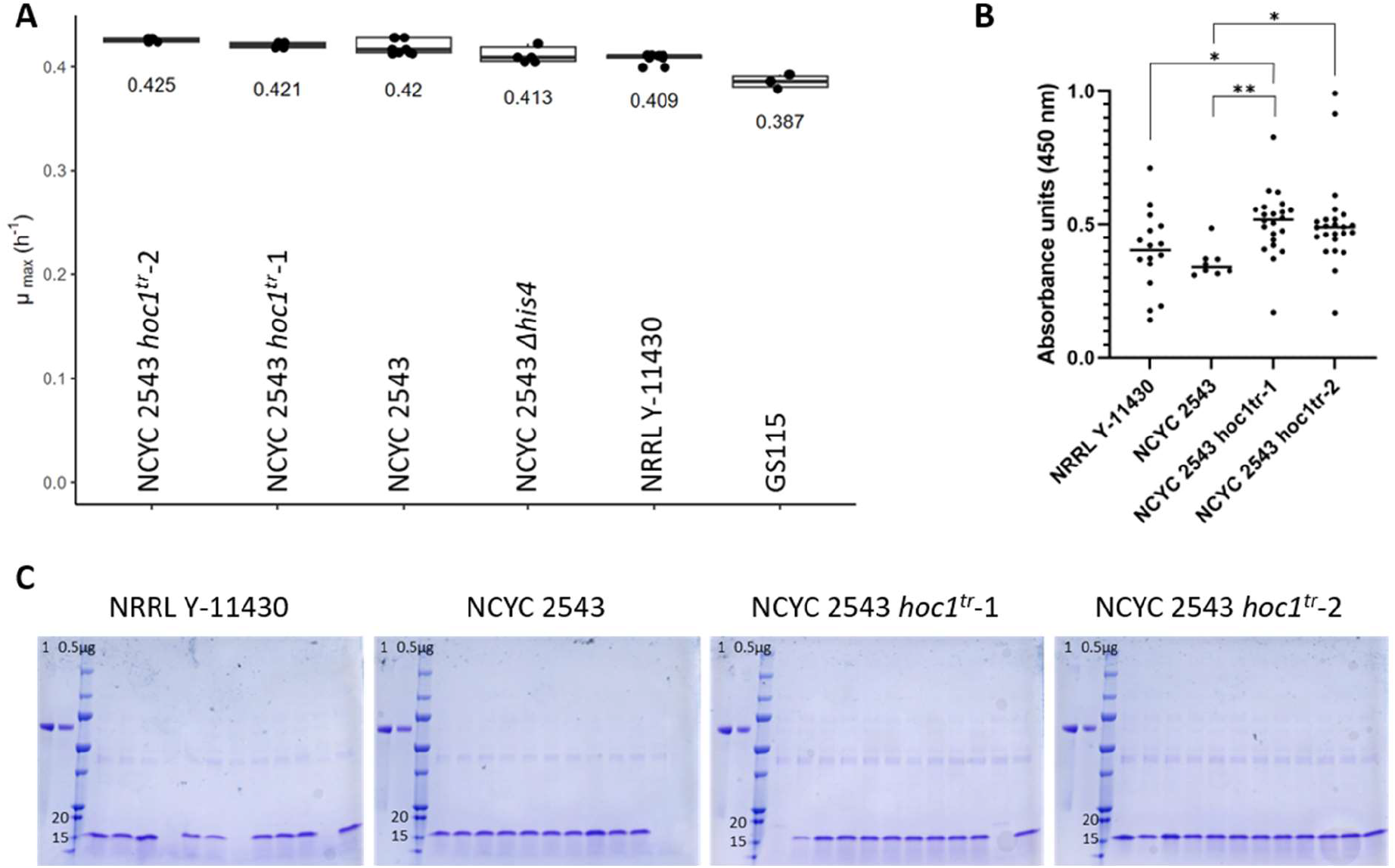
Overview of the strain performance of NRRL Y-11430, NCYC 2543 and the two NCYC 2543/OPENPichia hoc1^tr^ mutants. A. Comparison of the maximal growth rate of the four main strains, accompanied by GS115 and the NCYC 2543/OPENPichia his4. B. ELISA results of 24 clones of pGAP-based GBP expression in the different strains (no selection for single copies was done, wells were excluded when no expression was observed on SDS-PAGE, assuming these clones contain no expression cassette, or due to a technical issue during the ELISA procedure). Absorbance units were background corrected. All strains were compared in a Kruskal-Wallis omnibus test, followed by pairwise comparison corrected with Dunn’s multiple comparison procedure. Significance scores are annotated as follows, *: 0.002<p<0.03; **: 0.001<p<0.002; non-significant differences are not shown. C. SDS-PAGE analysis of the first 12 clones per transformation. Equal volumes of supernatant were loaded.

**Table 4.**
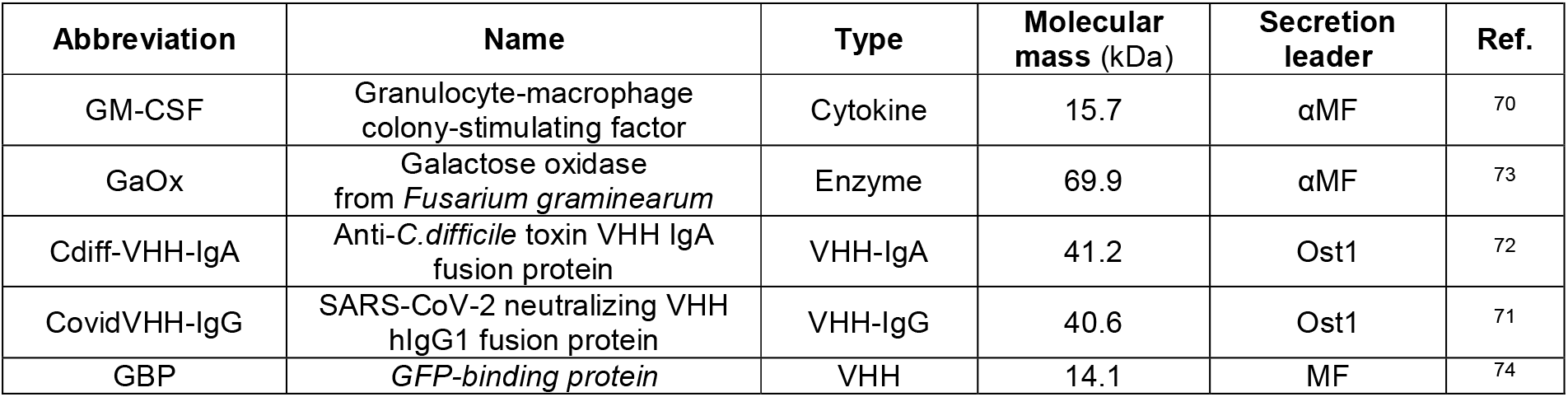
Details of the selected proteins used for the protein expression comparisons.

| Abbreviation | Name | Type | Molecular mass (kDa) | Secretion leader | Ref. |
| --- | --- | --- | --- | --- | --- |
| GM-CSF | Granulocyte-macrophage colony-stimulating factor | Cytokine | 15.7 | αMF | 70 |
| GaOx | Galactose oxidase from *Fusarium graminearum* | Enzyme | 69.9 | αMF | 73 |
| Cdiff-VHH-IgA | Anti-*C.difficile* toxin VHH IgA fusion protein | VHH-IgA | 41.2 | Ost1 | 72 |
| CovidVHH-IgG | SARS-CoV-2 neutralizing VHH hIgG1 fusion protein | VHH-IgG | 40.6 | Ost1 | 71 |
| GBP | *GFP-binding protein* | VHH | 14.1 | MF | 74 |

The presented data together with the growth and expression analysis, show that the NCYC 2543 background is at least as good as an expression host as the industrial NRRL Y-11430 strain (or its derivatives). With the discovery of the *HOC1* truncation as the basis for the higher transformability of NRRL Y-11430, and the finding that it can easily be accomplished using free-to-operate recombination-based genome engineering, also this last remaining handicap of the parental NCYC 2543 was removed. Indeed, as reported recently by Brady *et al*.^24^, the low transformability, which makes it laborious to find clones with multicopy integration of the expression vector, was the key reason for them to decide on continued use of NRRL Y-11430-based strains over more wild type strains. This problem is now solved.

As no difference between the two NCYC 2543 *HOC1*-mutant versions could be observed and given their excellent performance, we decided to continue all future *Pichia* work with the version *hoc1*^*tr*^*-1*, which has the smallest genomic scar downstream of the truncated *HOC1* coding sequence (i.e., insertion of a remaining Lox72 site).

### OPENPichia modular protein expression vector construction toolkit

Many scientists have used or still use commercial *K. phaffii* expression kits, which conveniently match expression strains and vectors. Indeed, constraints in vector choice are sometimes imposed by the properties of the chosen *K. phaffii* strain (e.g., selection markers, auxotrophies, methanol utilization phenotypes, etc). Although such kits are attractive to beginner users and are hence widespread within the scientific community, their conditions of sale are legally restrictive and forbid further distribution and reutilization of vector parts, let alone use in commercial production, which requires commercial licensing from the provider. Fortunately, issues with non-FTO DNA constructs can nowadays be avoided by *de novo* synthesis, combined with novel and fast cloning techniques which both emerged rapidly in the last decade^39^. However, establishing a new properly documented FTO genetic toolkit is still a relatively expensive and time-consuming occupation for most labs.

To enable scientists to express their genes of interest, a well-equipped genetic toolkit and corresponding cloning framework is provided to the community (Figure 8), and deposited at the BCCM plasmid collection. The cloning system that was chosen is modular, to enable maximum flexibility and based on Golden Gate cloning, similar to other toolkits^25,40–48^. These cloning systems have gained a lot of popularity as they are user-friendly and easily expandable, while also boasting high versatility in construct design. In essence, the strength of Golden Gate assembly is based on the use of Type IIS restriction endonucleases that cut outside their recognition sites, which allows users to flank DNA fragments of interest with customizable 4 nt overhangs. As such, a 4 nt overhang of one fragment can be made complementary to a 4 nt overhang of another fragment, and such compatible overhangs are ligated much more efficiently, enabling directional, multi-insert cloning in a single reaction. The MoClo system takes this concept a step further, as it standardizes Golden Gate assembly by designating *a priori* all DNA elements of a desired vector, which are typically referred to as ‘parts’, to a particular ‘part type’ (e.g., promoter, CDS, etc) and flanking each part type by unique 4 nt overhangs and Type IIS restriction sites^48^. In practice, parts are derived from PCR fragments or synthetic constructs, which are first subcloned in entry vectors, also known as ‘Level 0’ vectors (Figure 8). As such, a collection of sequence-verified Level 0 vectors is established and vectors of interest can then be assembled into expression vectors of interest, which are termed ‘Level 1’ vectors. By providing proper connector sequences with additional Type IIS restriction sites, the resulting expression vectors or Level 1 vectors can then be assembled again to obtain multigene or Level 2 vectors, which is the top level in the system’s hierarchy. In the current toolkit, all 4 nt overhangs were adopted to ensure a high degree of compatibility with existing yeast toolkits^25,41,45^ and to ensure a near 100% predicted ligation fidelity^49^. Since this toolkit is essentially derived from the *S. cerevisiae* MoClo system, it shares the restriction enzymes (BsmBI and BsaI), most of the 4 nt overhangs, and the number and design of the individual part types^41^. As such, the system is comprised of eight part types, of which Part 3 (Coding Sequence) and Part 4 (Terminator) can still be split up to allow additional modularity, for example to incorporate N- and C-terminal fusion partners for the protein of interest (Figure 8). An overview of the part types and the parts that are provided in this OPENPichia toolkit is included (Supplementary Figure 3). Part sequences are available in Supplementary Information and materials can be obtained from the BCCM GeneCorner plasmid collection^50^. The Material Transfer Agreement associated with it was custom-designed in collaboration with GeneCorner to allow for any use of resulting plasmids, including royalty-free commercial manufacturing.

**Figure 8.**
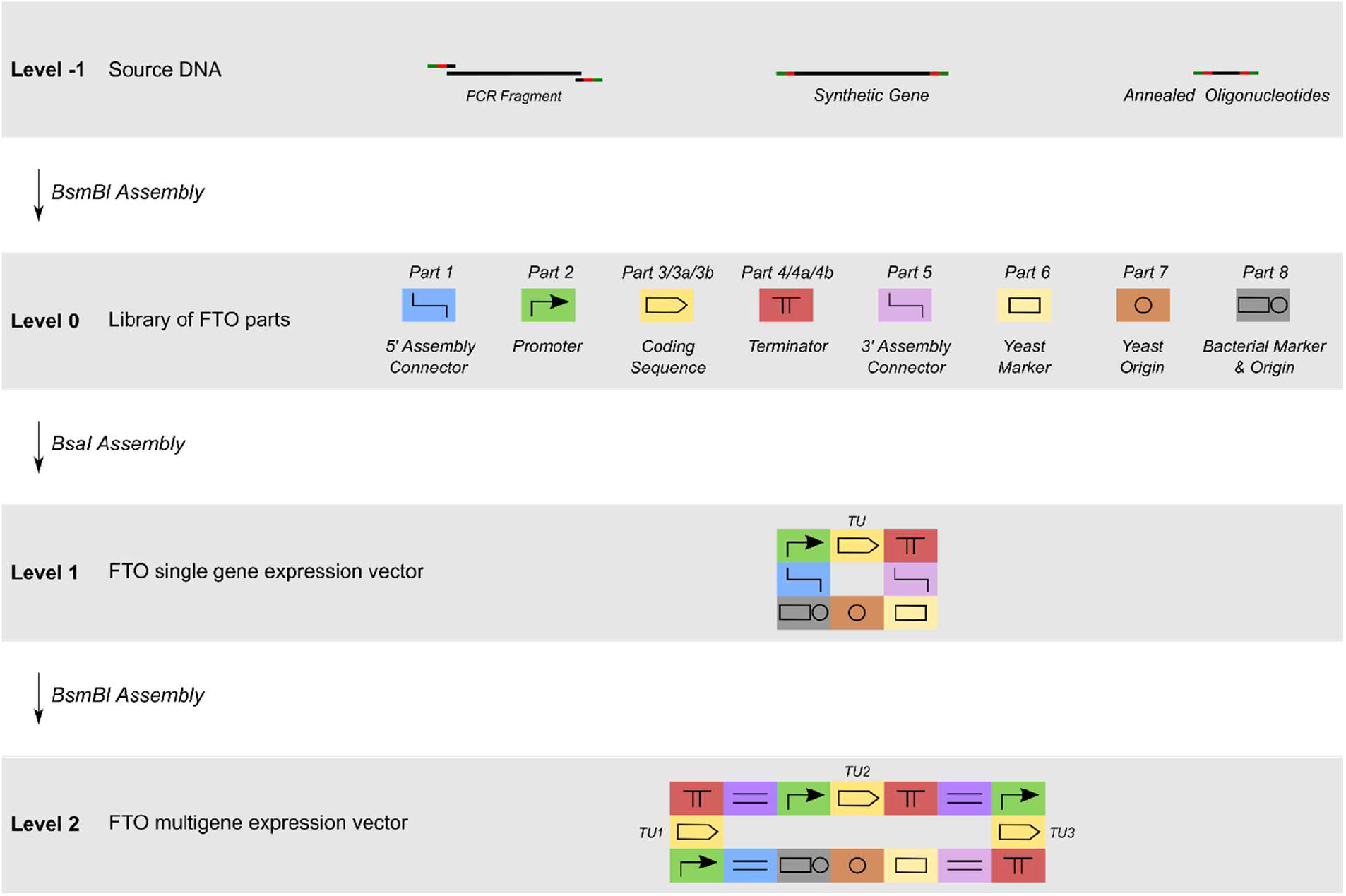
The modular cloning or MoClo principle. In a first phase, source DNA, such as PCR fragments, synthetic genes or annealed oligonucleotides are flanked with the proper Type IIS restriction sites and 4 nt overhangs, which are then accommodated in a Level 0 entry vector through BsmBI digest and T4 DNA ligation. Then, selected Level 0 vectors are assembled into a Level 1 expression vector by means of a BsaI digest and T4 DNA ligation. Finally, the system allows the assembly of multiple transcription units (promoter, CDS, terminator) from the individual Level 1 vectors, into a higher order Level 2 vector, in case the assembly connector sequences were properly selected during the assembly of the Level 1 vectors. Assembled assembly connectors are depicted with two horizontal lines and different shades of blue and purple. Note that the Part 3 Coding Sequence can be split up in a Part 3a and Part 3b Coding Sequence, to allow additional modularity. Likewise, the Part 4 Terminator can be split up in a Part 4a Coding Sequence and a Part 4b Terminator. Figure adapted from Lee et al^41^.

## Discussion

*Pichia pastoris (*formally known as *K. phaffii*) is an important protein production host in both academia and industry, but the most common industrially developed strains are currently distributed with restrictive MTAs, or not any longer. To facilitate academic and commercial host strain development for recombinant protein expression and easy distribution throughout the biotechnology community, alternative *K. phaffii* strains were investigated. We put forward an open-access wild type strain, i.e., NCYC 2543 or OPENPichia, as well as two derivatives: a histidine-auxotrophic strain and a *HOC1*-truncated strain with an improved transformation efficiency. Moreover, a compatible genetic engineering toolkit is made available, which contains all the necessary components for the expression of recombinant proteins and which can be easily expanded with more genetic parts to fit the researcher’s needs. This toolkit can be obtained with an open-usage MTA from the GeneCorner plasmid collection, but it can also be assembled from scratch, based on the sequences provided in the Supplementary Files 2.

It was shown that the proposed alternative OPENPichia strain (NCYC 2543) and its derivatives are almost identical to the common NRRL Y-11430 strain. Only a handful of mutations could be identified in our direct comparative genome analysis, of which only 4 are protein coding-altering (SNPs and indels). Additionally, OPENPichia does not contain the undesired killer plasmids and the strain shows the same maximum growth rate under the tested conditions. With respect to the protein production capacity, the data presented here demonstrate that small differences can occur between the *K. phaffii* type strain NCYC 2543/OPENPichia and NRRL Y-11430, but that there is no consistently better performing strain, considering the variety of proteins tested in this study. Previously, Brady et al^24^, performed a similar experiment where NRRL Y-11430 showed to have the highest protein expression level as compared to a series of other *K. phaffii* strains. However, none of the type strains from which NRRL Y-11430 directly derives were included in this study; instead, Y-12729 and Y-48124 (amongst others) were included, which are all members of Cluster 1 of their transcriptomics experiment. Y-7556 and YB-4290, which are type strains like NCYC 2543, are members of the transcriptomics Cluster 2, and would have allowed a better comparison. Due to the increased cell wall robustness and reduced transformation efficiencies of the type strains, they were excluded from the protein expression comparison in that study. Indeed, we also observed that the transformation efficiency of the NCYC 2543 strain is dramatically lower as compared to NRRL Y-11430. However, we could completely overcome this after our discovery by more in-depth genome resequencing that NRRL Y-11430 has a truncating frameshift mutation in the *HOC1* cell wall synthesis gene. We introduced the same frameshift mutations in *HOC1* of OPENPichia, resulting in an even improved transformation efficiency as compared to NRRL Y-11430. Hoc1p is part of one of two Golgi mannan polymerase complexes that in *S. cerevisiae* also contains Anp1p, Mnn9p, Mnn10p and Mnn11p^51^, and which mediates elongation of the α-1,6-mannan backbone that is initiated by the activity of Och1p. In *S. cerevisiae, HOC1* has synthetic positive genetic interactions with the *PKC1* pathway that mediates responses to cell wall stress, as well as genetic interactions with a multitude of genes involved in cell wall integrity, protein secretion, vesicular transport, all key pathways in the yeast cell wall maintenance^52^. Indeed, we observed a lower cell wall mannoprotein N-glycan mannosylphosphorylation density, which is a sensitive hallmark of cell wall stress. The truncated Hoc1p that is produced in NRRL Y-11430 vs. the type strain still has the alpha-helicoidal N-terminal luminal protein regions that type II Golgi proteins typically use to space their catalytic domains away from the membrane, and to interact with other Golgi proteins of similar topology. As Hoc1p forms part of such multi-protein complex, we hypothesize that the truncated *Hoc1* allele maintains assembly of this complex but lacks its own catalytic activity (see AlphaFold 2 model of the protein in Supplementary Figure 4). In this way, likely a milder phenotype is obtained than with full *HOC1* deletion, making the *hoc1*^*tr*^ strains grow equally well as the wild type.

Interestingly, under pGAP expression conditions, NRRL Y-11430 has somewhat more HCPs in its culture supernatant and grows to a lower cell culture density as compared to OPENPichia. We hypothesize that both observations are related and due to slightly increased cell lysis in NRRL Y-11430, which can have an impact on the need for additional purification steps. Whether the *hoc1*^*tr*^ mutation in OPENPichia has the same effect remains to be determined.

The *K. phaffii* strain that is proposed here as OPENPichia is one deposit of the type strain which is widely available in many culture collections in different countries (see Global Catalogue of Microorganisms)^53^. For instance, strain CBS 2612 also does not have killer plasmids and is identical but for a few drift mutations. If a *Pichia*-user is an end-user and wishes to merely manufacture a product in these type strains, multiple culture collections provide cost-effective licenses. However, for labs who are developing improved *Pichia* technology and implement these novel inventions in the strains, strains have to be distributable to other laboratories, and this is most often not allowed by the MTAs even of the public strain collections. NCYC should be commended for uniquely transparently providing cost-effective resale/redistribution as well as commercial manufacturing use licenses as part of standard culture collection practice, fulfilling an essential need for yeast-based technology developers.

More broadly, our study illustrates the need to build ‘generic’ biotechnological platforms after patents on these foundational inventions of our field expire, much in the same way as ‘generic/biosimilar’ medicines need to be developed to increase access to more affordable medicines. We have previously also accomplished this for the HEK293 cell lineage that is used for viral vector and vaccine manufacturing and hope that others will join us in such open science endeavours for other synthetic biology ‘chassis’ systems^54^. For now, we invite all *Pichia* researchers and users to contribute to this OPENPichia resource and make best use of it.

## Materials & Methods

### Strains and Media

The wild type *K. phaffii* strains NRRL YB-4290, NRRL Y-7556 and NRRL Y-11430 were obtained from the Agricultural Research Service (ARS, USA), CBS 2612 was obtained from Westerdijk Institute (Netherlands) and NCYC 2543 was obtained from the National Collection of Yeast Cultures (NCYC, UK). All mentioned strains were cultured and maintained on YPD or YPD agar.

All entry vectors and expression vectors were propagated and are available in the *E. coli* DH5α strain. MC1061 and MC1061λ strains were used successfully as well and generally showed higher transformation efficiency and easier green-white or red-white screening than was the case for DH5α. All *E. coli* strains were cultured and maintained on LB agar.

Antibiotics were used in the following concentrations for selection in *E. coli*: Zeocin 50 µg/ml, Nourseothricin 50 µg/ml, Hygromycin 50 µg/ml, Kanamycin 50 µg/ml, Chloramphenicol 50 µg/ml and Carbenicillin 50 µg/ml. Antibiotics were used in the following concentrations for selection in *Pichia*: Zeocin 100 µg/ml, Nourseothricin 100 µg/ml, Hygromycin 100 µg/ml, Geneticin 100 µg/ml, Blasticidin 100 µg/ml.

Several media were used: LB (1% Tryptone, 0.5% Yeast Extract, 0.5% NaCl), YPD (1% Yeast Extract, 2% Peptone, 2% D-glucose), YPG (1% Yeast Extract, 2% Peptone, 1% Glycerol), BMY (1% Yeast Extract, 2% Peptone, 1,34% YNB without amino acids, 100 mM Potassium phosphate buffer pH6), BMGY (BMY with 1% Glycerol), BMDY (BMY with 2% D-glucose), BMMY (BMY with 1% Methanol), and limiting glucose (1% Yeast Extract, 2% Peptone, 100mM Potassium phosphate buffer pH6, 50g/l Enpresso EnPump substrate, 5ml/l Enpresso EnPump enzyme solution (Enpresso GmbH, Germany)). For plates, 1.5% agar was added for LB media and 2% for YPD media; when Zeocin selection was used, media were set to pH7.5. All oligonucleotides and synthetic DNA fragments were ordered at Integrated DNA Technologies (IDT), Leuven, Belgium. All synthetic DNA fragments (gBlocks^®^ and Genes^®^) were designed and adapted for synthesis using the Codon Optimization Tool and the gBlocks Gene Fragments Entry Tool available at the website of IDT Europe.

### Illumina sequencing

The strains were cultured overnight in YPD medium and the genomic DNA was extracted using the Epicentre MasterPure™ Yeast DNA Purification Kit. Sample preparation (DNA fragmentation, adapter ligation, size selection and amplification) and next generation sequencing (5M 150bp paired end reads) was done by Eurofins, using Illumina technology. The reads were checked for quality using fastqc^55^, from which the %GC and number of reads was obtained. From the number of reads, the average overall coverage was calculated with the formula

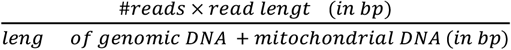

#### NGS analysis

The reads were trimmed using Trimmomatic^56^ to remove adapter, leading and trailing low quality bases (cut off quality 3), low quality reads (4-base sliding window quality below 15) and reads below 100 bp. Next, the reads were aligned to a reference and the mutations were identified using Breseq^27^ in consensus mode. As reference, the genome sequence published by Sturmberger *et. al*.^26^ was used. The reference sequences for killer plasmids and the mitochondrial DNA were obtained from Sturmberger *et. al*.^26^ and Brady *et. al*.^17^, respectively. The reported coverage depth was calculated by the Breseq algorithm. This is done by fitting a negative binomial distribution to the read coverage depth observed at unique reference positions. The mean of this binomial fit is used as the coverage depth. The killer plasmid copy number was estimated by comparing their coverage depth with the average of the four chromosomes. The coverage depth for each molecule was calculated as the mean of a binomial fit for the coverage depth for each reference position.

#### Phylogenetic tree

In order to make a phylogenetic tree, the sequencing data from this study was combined with the raw reads that were published before^24^ and also aligned as described above. From the predicted mutations of both datasets, a whole genome alignment was constructed from which a phylogenetic tree was calculated using the Mega X^57^ software package. A maximum likelihood algorithm was used with an HKY substitution matrix.

### Creation of the NCYC2543 *his4* strain

The NCYC2543 *his4* strain was generated using the split-marker method that was described previously by Heiss *et al*.^58^. The homology arms of the *HIS4* gene were selected from Näätsaari *et al*.^59^, and the reference genome of the CBS7435 strain. First, a construct containing the two homology arms with a floxed Nourseothricin acetyltransferase marker was created. Next two overlapping fragments containing one of the homologies and a part of the antibiotic marker were generated by PCR using Taq polymerase (Promega), which overlap for a length of 594 bp. These fragments were purified by a phenol chloroform precipitation. In brief, after adding an equal volume of phenol:chloroform:isoamyl alcohol (25:24:1), the solution was mixed, centrifuged (5 min at 12,000 g) and the liquid phase was isolated by decanting. To this, 1/10^th^ volume of 3 M sodium acetate pH 5.5 and 2 volumes of 100% ethanol was added and the sample was mixed and centrifuged (15 min at 12,000 g). Afterwards, the pellet, containing the amplified DNA, was isolated, washed with 70% ethanol, air-dried, and resuspended in water.

Both purified fragments were transformed into NCYC 2543 competent cells by electroporation and transformants were streaked to single clone onto YPD plates containing Nourseothricin and grown for 2 days at room temperature. The resulting clones were replica plated onto CSM-his plates for growth screening and grown for 2 days at room temperature. Strict non-growers were checked by colony PCR for replacement of the *HIS*4 gene with the antibiotic marker cassette.

The Nourseothricin acetyltransferase marker was finally removed by transient expression of a Cre-recombinase. This gene was cloned into a plasmid with an ARS^60^ and a Zeocin resistance cassette, which was then transformed in the *his4* strain. Transformants were incubated overnight on a YPD plate containing Zeocin and colonies were transferred to YPD plates without antibiotics. The removal of the antibiotic cassettes of the plasmid and *HIS4* knock-out was verified with replica plating on YPD containing the respective antibiotics and double-checked via colony PCR.

### Creation of the NCYC2543 *hoc1*^*tr*^ strains

The NCYC2543 *hoc1*^*tr*^ strains were generated using the split-marker method as described above. The left homology arm of the *HOC1* gene was chosen such as to contain about 1 kb upstream of the premature stop codon. To introduce the single nucleotide deletion, genomic DNA of NRRL Y-11430 instead of NCYC 2543 was used as the PCR template. The right homology arm was chosen as to contain about 1 kb downstream of the premature stop codon, also for this PCR, genomic DNA of NRRL Y-11430 was used, although NCYC 2543 would have also worked. The left and right homology arms were respectively fused by PCR to the first and last two thirds of the floxed Nourseothricin acetyltransferase marker. The PCR fragments were purified over gel and the DNA was recovered using the Wizard SV Gel and PCR Clean-Up System (Promega), according to the manufacturer’s instructions. Both purified fragments were transformed into NCYC 2543 competent cells by electroporation and transformants were streaked to single clone onto YPD plates containing Nourseothricin and grown for 2 days at room temperature. The resulting clones were screened using colony PCR using a forward primer annealing upstream of the left homology arm and a reverse primer annealing to the Nourseothricin selection marker. The Nourseothricin acetyltransferase marker was removed by transient expression of a Cre-recombinase as described above. The engineered *HOC1* locus was confirmed for both strategies by colony PCR and Sanger sequencing. The sequences for the PCR primers and split-marker cassettes can be found in Supplementary Tables 3 and 4.

### Growth Analysis

The different *Pichia* strains were cultured on YPD agar for 2 days, inoculated in triplicate into a 5 ml preculture of BMDY and grown overnight at 28 °C, shaking at 225 rpm. The optical density at 600 nm (OD600) of each culture was measured and 250 ml of BMDY was inoculated at a starting OD600 of 0.05. Samples of 1 ml were immediately isolated from each culture to measure the OD600 again. Then, samples of 1 ml were isolated every 2 h for 22 hours and again after 26 hours and 29 hours. All samples were diluted accordingly and measured within an OD600 range of 0.05 – 1.

### Recombinant protein expressions

The expression vectors were made using the MoClo toolkit, based on Golden Gate cloning as described in this paper. Briefly, the protein coding sequences were ordered synthetically with Part 3b type BsaI overhangs (NEB R3733) and cloned into the entry vector with BsmBI (NEB R0739). Next, expression vectors were made by assembly of the Level 0 parts.

The cloning procedure was as follows: 1 µl of T4 DNA Ligase (400 U; NEB M0202), 2 µl of T4 DNA Ligase Buffer (NEB M0202), 1 µl of Restriction Enzyme (20 U) were added to 20 fmoles of backbone (pPTK081 for entry vectors; any P8 backbone for destination vectors). An excess of insert (>1000 fmoles of PCR amplicon or synthetic gene; 10 pmoles of annealed oligonucleotides) was added for a BsmBI assembly, while equimolar amounts (20 fmoles) of each entry vector were added for a BsaI assembly. BsmBI assembly mixtures were incubated according to the following protocol: >25 cycles of 42 °C for 2 min (digest) and 16 °C for 5 min (ligation), followed by 60 °C for 10 min (final digest) and 80 °C for 10 min (heat inactivation step). BsaI assembly mixtures were incubated similarly, except that the digestion steps were performed at 37 °C.

*Pichia* electrocompetent cells were generated, using the lithium acetate method as described by Wu *et al*.^61^. In brief, precultures were inoculated in 5 ml YPD and grown overnight in an incubator at 28 °C and 250 rpm. The precultures were diluted and grown to an OD600 of approximately 1.5. 50 ml of the culture was isolated and the cells were harvested by centrifugation (1,519 g for 5 min at 4 °C), resuspended in 200 ml of a lithium acetate (LiAc)/dithiothreitol (DTT) solution (100 mM LiAc, 10 mM DTT, 0.6 M sorbitol, 10 mM Tris-HCl pH 7.5) and incubated for 30 minutes at 28 °C and 100 rpm. Next, the cells were collected by centrifugation (1,519 g for 5 min at 4 °C), washed two times with 1 M ice-cold sorbitol and finally resuspended in 1.875 ml of 1 M ice-cold sorbitol. 0.5 to 1 µg of DNA was added to aliquots of 80 µl and electroshocked (1.5kV, 200Ω, 25µF). Immediately, 1 ml of 1 M sorbitol was added and the suspension was incubated at 28 °C for 2-5 h. Next, the cells were plated on YPD agar containing the appropriate antibiotic, and colonies were isolated after 2 days of incubation at 30 °C.

To be able to compare expression, only colonies with single copy integration of the construct were selected. The copy number was determined by quantitative PCR on a Lightcycler 480 (Roche) using primers that bind pAOX1 and pGAP. The genes *OCH1* and *ALG9* were used as reference. Genomic DNA (gDNA) of NCYC 2543 was included as a single copy positive control. A single copy plasmid integration will yield one additional copy and more than two copies would be the result of multiple plasmid integrations. Amplification efficiencies were determined using serial dilutions of gDNA samples. Reactions were set up in 10 μl with final concentrations of 300 nM forward primer, 300 nM reverse primer, 1x SensiFast SYBR no-rox mastermix (Bioline), 10 ng gDNA and the following cycling conditions: 3 min at 95 °C, followed by 45 cycles of 95 °C for 3 s, 60 °C for 30 s at ramp rate 2.5 °C/s, 72 °C for 1 s, ending with 0.11 °C/s from 65 °C to 95 °C for melting curve determination (5 acquisitions/s). Copy numbers were calculated using the ΔΔCt method^62^.

The different strains expressing the recombinant proteins, were cultured on YPD agar plates for 2 days, inoculated in triplicate into a 5 ml preculture of BMDY and grown overnight at 28 °C, shaking at 225 rpm. Next, the cultures for pAOX1-driven expression, were inoculated in BMDY, grown for 24 h, subsequently transferred to BMMY and incubated for 48 h. After 24 h in BMMY, an extra 1% of methanol was added. The cultures for pGAP-driven expression, were instead inoculated in limiting glucose medium and incubated for 48 h. Then, optical density at 600nm was measured for all cultures and the supernatant was collected by centrifuging (2,500 g for 5 min). The samples were incubated with EndoH (produced in-house) to remove N-glycans and analysed by SDS-PAGE.

### ELISA-based quantification of GBP

Nunc MaxiSorp™ 96-well plates were coated with 75 ng/well of penta-His antibody in PBS solution (Qiagen, 34660) and incubated overnight at 4 °C. Plates were washed three times with 200 µl/well of wash buffer (PBS + 0.05% Tween-20) and any residual liquid was removed. Plates were blocked with 100 µl/well Reagent Diluent (1% Probumin (Millipore, 82-045-1) in PBS pH 7.2) for 2 h. Plates were washed three times with 200 µl/well of wash buffer and residual liquid was removed. Dilutions of the yeast supernatant were prepared in 96-deepwell plates and 100 µl of a 100,000-fold dilution was applied to the plates and incubated for 1 hour while shaking gently on a table top plate shaker. Plates were washed three times with 200 µl/well of wash buffer and residual liquid was removed. Plates were provided with 100 µl of a 250 ng/ml MonoRab™ Rabbit Anti-Camelid VHH Antibody coupled to HRP in Reagent Diluent and incubated for 1 hour while shaking gently on a table top plate shaker. Plates were washed three times with 200 µl/well of wash buffer and residual liquid was removed. TMB substrate was prepared according to the manufacturer’s instructions (BD OptEIA™) and 100 µl/well was applied to the plate before incubating for 10 min. Next, 50 µl of stop solution (2N H2SO4) was added to each well and the plate was read at 450 nm by a plate reader. Absorbance units were background corrected. All strains were compared in a Kruskal-Wallis omnibus test, followed by pairwise comparison corrected with Dunn’s multiple comparison procedure.

### Transformation efficiency testing

Competent cells were prepared using the lithium acetate method as described above. 200 ng of linearized plasmid was transformed to each strain and several dilutions of the transformation mix were plated on either non-selective YPD agar or YPD agar containing 100 µg/ml Zeocin. For each transformation, colonies were counted from the plates where clear individual colonies could be observed after 2 days at 30 °C incubation. Both the selective and non-selective plates were counted to correct for a potential difference in the number of competent cells per transformation.

A linear model (estimated using ordinary least squares) in the statistical software R was fitted^63^. As the outcome variable, the log-transformed, normalized transformation efficiency (natural log of the number of transformants per million clones) and as predictor variables the strain and promotor type, including an interaction effect were used. The model explains a statistically significant and substantial proportion of variance (R^2^=0.94, F(7,38)=81.33, p<.001, adj.R^2^=0.93). Model-predicted group means with 95% confidence intervals were obtained using the ggeffects package with heteroscedasticity-consistent variance estimators from the sandwich package (vcovHC, type HC0)^64,65^. Further, we defined specific contrasts using the multcomp package^66^, again with heteroscedasticity-consistent variance estimators to obtain multiple-comparison corrected estimates for the ratios of transformation efficiencies between the different strains and using different plasmids.

### DSA-FACE-based glycan analysis of the cell wall mannoproteins

Strains were inoculated in YPD or YPG, from their respective precultures, at an OD600 of 0.05 and grown overnight at 28 °C and 200 rpm. The next day, 500 OD units per strain were pelleted (10 min at 1,500 g) and the mannoproteins were isolated, as follows. The pellets were washed three times with Milli-Q, after which 20 mM of citrate buffer pH6.6 was added at 1 ml per 150 µg of wet cell weight. The resuspended cells were autoclaved for 1.5 hours at 120 °C in cryovials and then centrifuged for 10 min at 16,000 g. To the supernatant fractions, 3 volumes of ice-cold methanol were added and the vials were incubated for 15 min at 20 °C. The mannoproteins were spun down for 10 min at 16,000 g and the pellets were left to dry until transparent. The pellets were resuspended in 50 µl RCM buffer (8 M Urea, 360 mM Tris-HCl pH 8.6, 3.2 mM EDTA) and stored at 4 °C until further analysis.

The N-linked oligosaccharides were prepared from the purified mannoproteins upon blotting to PVDF membrane in the wells of 96-well plate membrane plates, and were analysed by capillary electrophoresis with laser-induced fluorescence detection (CE-LIF) using an ABI 3130 capillary DNA sequencer as described previously^36^.

### Alcian blue assay

The assay was performed as described previously^37^, with adaptations. Briefly, Alcian blue was prepared in 0.02 N HCl at a concentration of 63 µg/ml and the solution was centrifuged to remove insoluble precipitates. An overnight culture of each strain was grown in YPD at 28 °C and 200 rpm. The next day, the cells were pelleted and the supernatant was removed. The cells were washed with 0.02 N HCl and the pellet was resuspended again in 0.02 N HCl to an OD600 of 10 OD/ml. 100 µl (1 OD600) of cells was transferred to a 96-V-bottom plate to which 100 µl of the Alcian blue solution was added. After 15 min of incubation at room temperature, the plate was centrifuged for 15 min at 3,220 g, after which the pellets were visually checked.

### Congo red and Calcofluor white test

The test was performed as described elsewhere^67^, with slight adaptations. Briefly, the strains were grown overnight in BMGY. The next day, dilutions were made in order to obtain 10E5 to 10E1 cells in 5 µl BMGY. 5 µl drops were spotted on the different plates and the plates were incubated for 3 days at 30 °C. Congo red (Sigma, C6767) and Calcofluor white (Fluka, 18909) were present at final concentrations of 75 µg/ml and 10 µg/ml, respectively.

### Electron microscopy

#### Transmission electron microscopy

The strains were cultivated in BMGY at 28 °C and 200 rpm, overnight. High Pressure Freezing, as described previously^68^, was carried out in a high-pressure freezer (Leica EM ICE; Leica Microsystems, Vienna, Austria). Cells were pelleted and frozen as a paste in 150 µm cupper carriers. HPF was followed by Quick Freeze Substitution as described previously^69^. Briefly, carriers were placed on top of the frozen FS solution inside a cryovial containing 1% ddH2O, 1% OsO4 and 0.5% glutaraldehyde in dried acetone. After reaching 4 °C for 30 min, samples were infiltrated stepwise over three days at 0-4 °C in Spurr’s resin and embedded in capsules. The polymerization was performed at 70 °C for 16 h. Ultrathin sections of a gold interference colour were cut using an ultra-microtome (Leica EM UC6), followed by a post-staining in a Leica EM AC20 for 40 min in uranyl acetate at 20 °C and for 10 min in lead stain at 20 °C. Sections were collected on formvar-coated copper slot grids. Grids were viewed with a JEM-1400Plus transmission electron microscope (JEOL, Tokyo, Japan) operating at 60 kV.

#### Scanning electron microscopy

The strains were cultivated in BMGY at 28 °C and 200 rpm, overnight. Cells were fixed overnight in 1.5% Paraformaldehyde, 3% Glutaraldehyde in 0.05 M Na-Cacodylate buffer pH7.4. The fixed cells were centrifuged for 2 min at 1,000 g between each following step. First the cells were washed 3 times with 0.1 M Na-Cacodylate buffer pH7.4 and then incubated for 30 min in 2% OsO4 in 0.1 M Na-Cacodylate pH7.4. Osmicated samples were washed 3 times with Milli-Q, prior to a stepwise ethanol dehydration (50%, 70%, 90%, 2 × 100%). Samples were incubated twice in hexamethyldisilazane (HMDS) solution (Sigma-Aldrich), as a final dehydration step, after which they were spotted on silicon grids (Ted Pella) and air-dried overnight at room temperature. Samples were next coated with 5 nm Platinum (Pt) in a Quorum Q 150T ES sputter coater (Quorum Technologies) and placed in a Gemini 2 Cross beam 540 microscope from Zeiss for SEM imaging at 1.50 kV using a SE2 detector.

## Associated content

- All plasmids from the modular cloning kit are available at the Belgian Co-ordinated Collections of Micro-organisms (BCCM)/GeneCorner Plasmid Collection (http://bccm.belspo.be/about-us/bccm-genecorner).
- All raw reads of the genomes sequenced in this study have been submitted to NCBI and can be found under the following accession numbers: NRRL Y-11430 (SAMN32067769), NRRL YB-4290 (SAMN32067770), NRRL Y-7556 (SAMN32067773), NCYC 2543 (SAMN32067771), CBS 2612 (SAMN32067772).
- Supplementary Figure 1: Summary of the end-ODs of the pGAP- and pAOX1-based cultivations at harvest.
- Supplementary Figure 2: DSA-FACE profiles of the cell wall mannoproteins of the different strains grown on YPD or YPG.
- Supplementary Figure 3: Overview of the available elements for the different parts of the MoClo toolbox.
- Supplementary Figure 4: AlphaFold 2 models of the type strain Hoc1p and of the NRRL Y-11430 lineage derived strain.
- Supplementary Table 1: Overview of the NGS results, including the number of reads, GC% and average overall coverage.
- Supplementary Table 2: Proportion of NGS reads mapping to different molecules of the reference genome, mitochondrial DNA and killer plasmids.
- Supplementary Table 3: List of oligonucleotides that were used as primers for PCR, cPCR or sequencing.
- Supplementary Table 4. Sequences of the split-marker fragments used to generate the two *HOC1* mutants.
- Supplementary File 1: GenBank files of the expression constructs used in the study.
- Supplementary File 2: FASTA files of the available MoClo Parts.

## Author Information

This work was originally conceived and initiated by DVH, KV and NC. DVH, RV, KV, SV, EW, BVM, HE, DF, HG, CL, GM, LM, JN, CR, LVS, and KC performed experiments and contributed to data analysis and/or results presentation. RDR, MDB and PB performed the electron microscopy. DVH, RV, KC and NC cowrote the manuscript, while KC and NC supervised the work.

## Acknowledgements

DVH was supported by a Baekeland mandate of VLAIO (Flanders Innovation & Entrepreneurship fund) in collaboration with Inbiose NV, and is now an employee of Inbiose NV. RV was supported by a Strategic Basic Research fellowship from the Fund for Scientific Research and otherwise supported by Ghent University and is now an employee of Those Vegan Cowboys. KV was a VIB post-doctoral fellow and is now employee of Inbiose NV. SV and EW are employees of the Ghent University. HE is supported by a Fundamental Research fellowship of the Fund for Scientific Research Flanders (FWO). HG was a post-doctoral fellow funded by the Ghent University and VIB and is now an employee of Eurofins. CL was supported by a Strategic Basic Research fellowship of the Fund for Scientific Research Flanders (FWO) and is now an employee of Exevir. LM and DF are staff scientists of the VIB Center for Medical Biotechnology. GM was supported by VIB and now works at Animab. JN, CR and BVM are supported by Strategic Basic Research fellowships of the Fund for Scientific Research Flanders (FWO). LVS is a VIB post-doctoral fellow and supported by grants from the Industrial Research Fund of Ghent University, VLAIO and the European Commission (HERA-Pilot). RDR, MDB and PB are employees of VIB. KC is supported by an Innovation Mandate of VLAIO. Research in the Callewaert lab is supported by grants from UGent, the Fund for Scientific Research Flanders (FWO) and core resources from VIB. We want to thank Joeri Beauprez for valuable discussions; and Erhan Çıtak, Merve Arslan, Annelies Van Hecke and Simon Devos for assistance with some of the experiments.

## Notes

### Competing Interest Statement

The authors have declared no competing interest.

