## Supplementary Information for "OPENPichia: building a free-to-operate *Komagataella phaffii* protein expression toolkit"

### Supplemental material

#### Tables

**Supplementary Table 1. Overview of the next generation sequencing results.** For each sequenced strain, the total number of reads, %GC and average overall coverage are reported.

| Strain Designation | Number of reads | %GC | Average overall coverage |
|---|---|---|---|
| NRRL Y-11430 | 11883858 | 37 | 191 |
| NRRL YB-4290 | 16291010 | 31 | 261 |
| NCYC 2543 | 14748310 | 37 | 236 |
| CBS 2612 | 13586250 | 39 | 218 |
| NRRL Y-7556 | 13886574 | 32 | 223 |

**Supplementary Table 2. Proportion of NGS reads mapping to different molecules of the reference genome, mitochondrial DNA and killer plasmids.**

| Strain Designation | Reads mapped to | | |
|---|---|---|---|
| | Genome (%) | Mitochondrial DNA (%) | Killer plasmids (%) |
| NRRL Y-11430 | 79.6 | 17.7 | 2.7 |
| NRRL YB-4290 | 45.1 | 47.3 | 7.7 |
| NCYC 2543 | 78.5 | 21.6 | 0 |
| CBS 2612 | 89.4 | 10.7 | 0 |
| NRRL Y-7556 | 52.4 | 38.7 | 9 |

**Supplementary Table 3. List of oligonucleotides that were used as primers for PCR, cPCR or sequencing.**

| Primer name | Sequence (5' → 3') | Used for |
|---|---|---|
| OCH1_Fw | CCTCTGATAGTTCCTTTCCG | Fw primer in qPCR (copy number determination) for reference gene *OCH1* |
| OCH1_Rv | AAGACTTCTGGTACACGTTC | Rev primer in qPCR (copy number determination) for reference gene *OCH1* |
| ALG9_Fw | CTTTAGTGGGATGTTACCAG | Fw primer in qPCR (copy number determination) for reference gene *ALG9* |
| ALG9_Rv | CAACGTAAAAATCACACTCC | Rev primer in qPCR (copy number determination) for reference gene *ALG9* |
| AOX1p_Fw | CCATCCGACATCCACAGGTC | Fw primer for *AOX1* promoter copy number determination |
| AOX1p_Rv | TCGATGGCAAAAGTGGGTGT | Fw primer for *AOX1* promoter copy number determination |
| 72_HOC1_1a_Fw | GAGTGTCGCACGTCTCGG | Fw primer left homology arm *HOC1* (strategy 1&2) |
| 73_HOC1_1b_Rev | ATGCTATACGAACGGTATCAAGATTGGGTGACCAGGC | Rev primer left homology arm *HOC1* – overlap with Nour selection marker (strategy 1&2) |
| 74_HOC1_2a_Fw | CATTATACGAACGGTAGAGAATTAATCGCCAAAATCACAGAGG | Fw primer right homology arm *HOC1* – overlap with Nour selection marker (strategy 1) |

| | | |
|---|---|---|
| 75_HOC1_2b_Rev | GGATTCAATCCCGGATTTGTCG | Rev primer right homology arm *HOC1* (strategy 1&2) |
| 76_Lox71_3a_Fw | CACCCAATCTTGATACCGTTCGTATAGCATACATTATACG | Fw primer first two thirds of Nour selection marker – overlap with left homology arm *HOC1* (strategy 1&2) |
| 77_2/3Nour_3b_Rev | CAACGGTCAACCTTCTATTCC | Rev primer first two thirds of Nour selection marker (strategy 1&2) |
| 78_2/3Nour_4a_Fw | CTCGAAACCGCGACTTC | Fw primer second two thirds of Nour selection marker (strategy 1&2) |
| 79_Lox66_4b_Rev | GCGATTAATTCTCTACCGTTCGTATAATGTATGCTATACG | Rev primer second two thirds of Nour selection marker – overlap with right homology arm *HOC1* (strategy 1) |
| 82_HOC1_2c_Fw | CATTATACGAACGGTAGCAATGGACGGGTACTGG | Fw primer right homology arm *HOC1* – overlap with Nour selection marker (strategy 2) |
| 83_Lox66_4c_Rev | CCGTCCATTGCTACCGTTCGTATAATGTATGCTATACG | Rev primer second two thirds of Nour selection marker – overlap with right homology arm *HOC1* (strategy 2) |
| 11_US_HOC1_Fw | GGGGTTCGATAATCAAACTTGCCTTGAAGG | Colony PCR primer upstream of left homology arm of *HOC1* |
| 12_DS_HOC1_Rev | CACCGATACCCAGACCGTATCAACCAC | Colony PCR primer downstream of right homology arm of *HOC1* |

**Supplementary Table 4. Sequences of the split-marker fragments used to generate the two *HOC1* mutants.**

| Fragment | Sequence (5' → 3') |
|---|---|
| *HOC1* left fragment – strategy 1&2 | gagtgtcgcacgtctcggtatttttttacttgtcgattctctcttactgtccaagatgtacgaaagatgtgaagtttctttgt<br>tggcgattttcttgcagcgcgatcgcggtcggtatgatttcccttcagaccaaaaacagatatagatattcaattgaaactcg<br>cccttccaacaccctgaacaccacccccctaatgcaacaacagctgtttagaaaagtaatttggctcacggttggactcataac<br>ggtgatattggtaataataaagatttcatccagcaagtccaccgccacagaccttcagaaagtattgaagaatgccaatattt<br>taccgcaggacgtgatcaattacaatagcagaaaagtaacggatgaactcgcttcaaagctggacgagattcaaaagaagtat<br>ctctcgaagcaagatgatagaattagcaagctcgaagctgaacgggcagatctactggaacaggttagatttctaaggaaccc<br>ccctgcaggatcaagtttaagagaaaaattggcttatctgtttccttataatgaaaacggcaaattccctgcttatatatggc<br>aaacatggaagtatggcttgaatgacgatcggtttggagaaaagttcaaagaaggcgaaactcagtgggcttcgaagaatcct<br>ggtttcgttcatgagttgtttaacgatgatacttccggtgtgtttattcaccatctgtatatcaatgttccagaagtgatcaa<br>agcatacgagctgcttcccaacataatcttgaaaatggacttcttcagatatttggttttatacgccaagggaggtgtctatg<br>cagacgttgatactatgcctcttcagcctgtaccaaactggattcctgaaaatgtctccccaaaaagcatcgggatgatcatt<br>ggaatacaaaacgatgctaacaatccagattggaaaaagattacgtacatcgtctacaattttccaattggtgtattcaagcg<br>aagcctggtcacccaatcttgataccgttcgtatagcatacattatacgaagttatatccttcagtaatgtcttgtttctttt<br>gttgcagtggtgagccattttgacttcgtgaaagtttctttagaatagttgtttccagaggccaaacattccacccgtagtaa<br>agtgcaagcgtaggagaaccaagactggcataaatcaggtataagtgtcgagcactggcaggtgatcttctgaaagtttctac<br>tagcagataagatccagtagtcatgcatatggcaacaatgtaccgtgtggatctaagaacgcgtcctactaaccttcgcattc<br>gttggtccagtttgttgttatcgatcaacgtgacaaggttgtcgattccgcgtaagcatgcatacccaaggacgcctgttgca<br>attccaagtgagccagttccaacaatctttgtaatattagagcacttcattgtgttgcgcttgaaagtaaaatgcgaacaaat<br>taagagataatctcgaaaccgcgacttcaaacgccaatatgatgtgcggcacacaataagcgttcatatccgctgggtgactt<br>tctcgctttaaaaaattatccgaaaaaatttctagagtgttgttactttatacttccggctcgtataatacgacaaggtgta<br>aggaggactaaaccatgggtactaccttagatgatacagcctacagatacagaacatcagtccctggtgatgctgaagcaatt<br>gaggctttagacggttcattcaccaccgacaccgtctttagagtaaccgccaccggtgatggatttaccttaagagaagtccc<br>agtcgaccctccattaactaaagtctttccagatgatgaatctgatgacgaaagcgacgacggagaagatggtgacccagatt<br>caagaactttcgtagcatacggtgatgacggtgatttggctggttttgtagtcgtttcttattcaggttggaatagaaggttg<br>accgttg |
| *HOC1* right fragment – strategy 1 | ctcgaaaccgcgacttcaaacgccaatatgatgtgcggcacacaataagcgttcatatccgctgggtgactttctcgctttaa<br>aaaattatccgaaaaaatttctagagtgttgttactttatacttccggctcgtataatacgacaaggtgtaaggaggactaa<br>accatgggtactaccttagatgatacagcctacagatacagaacatcagtccctggtgatgctgaagcaattgaggctttaga<br>cggttcattcaccaccgacaccgtctttagagtaaccgccaccggtgatggatttaccttaagagaagtcccagtcgaccctc<br>cattaactaaagtctttccagatgatgaatctgatgacgaaagcgacgacggagaagatggtgacccagattcaagaactttc<br>gtagcatacggtgatgacggtgatttggctggttttgtagtcgtttcttattcaggttggaatagaaggttgaccgttgaaga<br>tatagaagtcgccccagagcatagaggtcatggtgtaggaagagctttgatgggtttggctacagaatttgcaagagagagag<br>gagccggtcatttatggttagaagttactaatgttaacgcccctgctatccatgcttatagaagaatgggtttcacattatgt<br>ggtttagatactgctttatatgatggaacagcatctgacggtgaacaggccttgtatatgtctatgccttgcccttaagccga |

| | |
|---|---|
| | gtaactgacaataaaaagattcttgttttcaagaacttgtcatttgtatagtttttttatattgtagttgttctattttaatc<br>aaatgttagcgtgatttatattttttttcgcctcgacatcatctgcccagatgcgaagttaagtgcgcagaaagtaatatcat<br>gcgtcaatcgtatgtgaatgctggtcgctatactgataacttcgtatagcatacattatacgaacggtagagaattaatcgcc<br>aaaatcacagaggatacactgcaacgagccgagtcaaactcactggaactagctgacattagcgaagaaggcggcctgtctga<br>taagaatttgtccattatgcaatggacgggtactggtatttttacagatgccatatttacctattttaatgactacattcaaa<br>gtagtatctataccaaagttacttggaaagaattctccaaattgagaaagcccaagcttgtcagtgatgtattggtactgccg<br>attatcagcttctcggccggtgcaggtagtggaaaatcgactgaactgaacgatcccttagcattcgtacaacattattttga<br>aagattacataacgacaaccactaagggtcagaaccatttagattgtctggatctatcattatggccttgtttatagacaaag<br>aattgtatcctggactgaagggaagtttatagagtaatacccctctgacaaccaactcgaatgggtattttgagataatttcc<br>catatattattgtcttcactgggagacttacttcttctttgtagttccggtatcggcagcaaacttatggtggtcacatcttc<br>caaacttctcggtatagtagaactatttgtggttgatacggtctgggtatcggtgataatgcttgtaataatggattttgtgg<br>gtgtaactgaagctggtgtattagtggtagtggtagtggtggttgttactgtgctatctgtaggttctgggagtggtttctga<br>gccgggtttaaaattttgatctctgtggtatcttcgtctggatttcgactaaaaaatccgccttgaaatacgagtggaacagg<br>taaacgattgatggagcccagaagcactgtgtccaaggaagattcatcaaaagtgaaaggagtgccatcttcgtttccatcgt<br>taagatgagaacttttggcaaaaaaatcaatcagaacgtcatctatttctatggcgacaaatccgggattgaatcc |
| *HOC1* right fragment – strategy 2 | ctcgaaaccgcgacttcaaacgccaatatgatgtgcggcacacaataagcgttcatatccgctgggtgactttctcgctttaa<br>aaaattatccgaaaaaatttctagagtgttgttactttatacttccggctcgtataatacgacaaggtgtaaggaggactaa<br>accatgggtactaccttagatgatacagcctacagatacagaacatcagtccctggtgatgctgaagcaattgaggctttaga<br>cggttcattcaccaccgacaccgtctttagagtaaccgccaccggtgatggatttaccttaagagaagtcccagtcgaccctc<br>cattaactaaagtctttccagatgatgaatctgatgacgaaagcgacgacggagaagatggtgacccagattcaagaactttc<br>gtagcatacggtgatgacggtgatttggctggttttgtagtcgtttcttattcaggttggaatagaaggttgaccgttgaaga<br>tatagaagtcgccccagagcatagaggtcatggtgtaggaagagctttgatgggtttggctacagaatttgcaagagagagag<br>gagccggtcatttatggttagaagttactaatgttaacgcccctgctatccatgcttatagaagaatgggtttcacattatgt<br>ggtttagatactgctttatatgatggaacagcatctgacggtgaacaggccttgtatatgtctatgccttgcccttaagccga<br>gtaactgacaataaaaagattcttgttttcaagaacttgtcatttgtatagtttttttatattgtagttgttctattttaatc<br>aaatgttagcgtgatttatattttttttcgcctcgacatcatctgcccagatgcgaagttaagtgcgcagaaagtaatatcat<br>gcgtcaatcgtatgtgaatgctggtcgctatactgataacttcgtatagcatacattatacgaacggtagcaatggacgggta<br>ctggtatttttacagatgccatatttacctattttaatgactacattcaaagtagtatctataccaaagttacttggaaagaa<br>ttctccaaattgagaaagcccaagcttgtcagtgatgtattggtactgccgattatcagcttctcggccggtgcaggtagtgg<br>aaaatcgactgaactgaacgatcccttagcattcgtacaacattattttgaaagattacataacgacaaccactaagggtcag<br>aaccatttagattgtctggatctatcattatggccttgtttatagacaaagaattgtatcctggactgaagggaagtttatag<br>agtaatacccctctgacaaccaactcgaatgggtattttgagataatttcccatatattattgtcttcactgggagacttact<br>tcttctttgtagttccggtatcggcagcaaacttatggtggtcacatcttccaaacttctcggtatagtagaactatttgtgg<br>ttgatacggtctgggtatcggtgataatgcttgtaataatggattttgtgggtgtaactgaagctggtgtattagtggtagtg<br>gtagtggtggttgttactgtgctatctgtaggttctgggagtggtttctgagccgggtttaaaattttgatctctgtggtatc<br>ttcgtctggatttcgactaaaaaatccgccttgaaatacgagtggaacaggtaaacgattgatggagcccagaagcactgtgt<br>ccaaggaagattcatcaaaagtgaaaggagtgccatcttcgtttccatcgttaagatgagaacttttggcaaaaaaatcaatc<br>agaacgtcatctatttctatggcgacaaatccgggattgaatcc |

### Figures

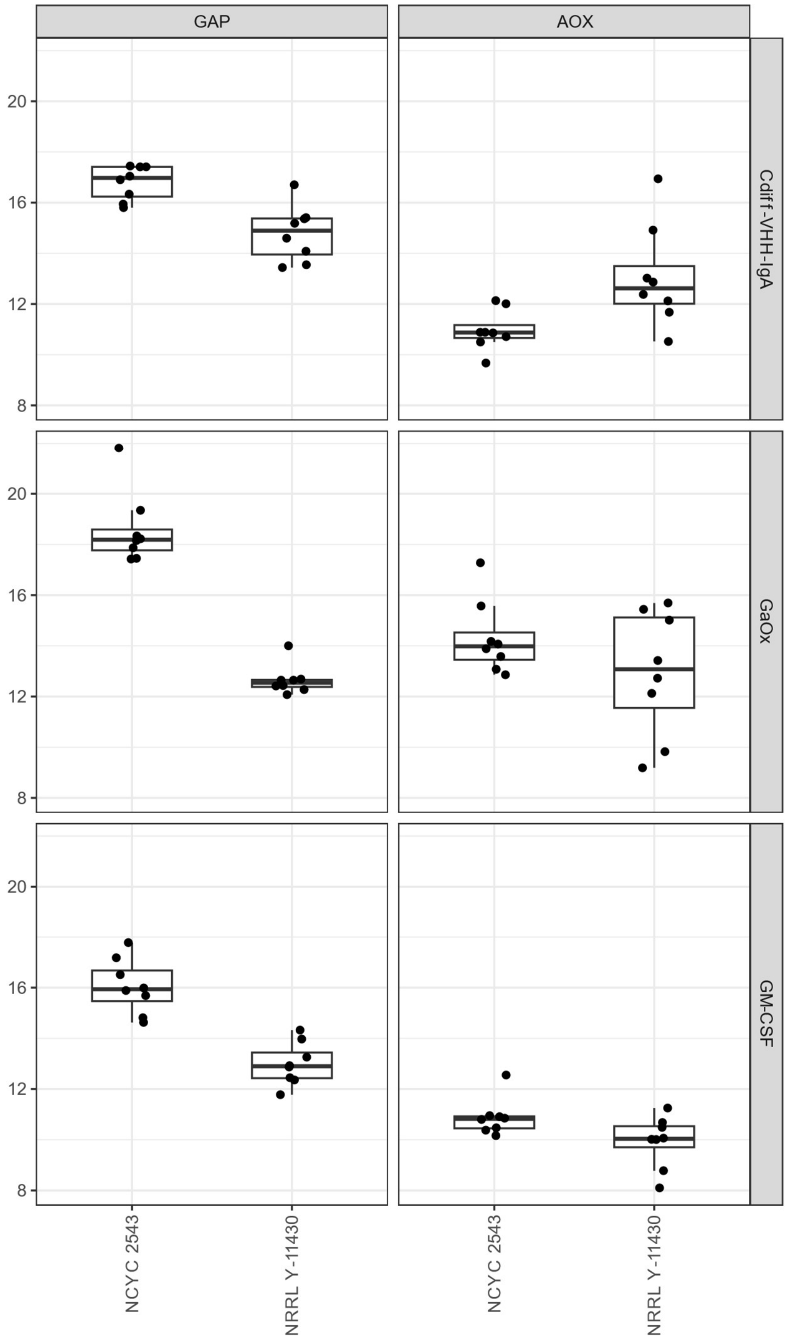


**Supplementary Figure 1. Summary of the end-ODs of the pGAP- and pAOX1-based cultivations at harvest.** For both strains, NCYC 2543 and NRRL Y-11430, the end-ODs for the three model protein cultivations are depicted.

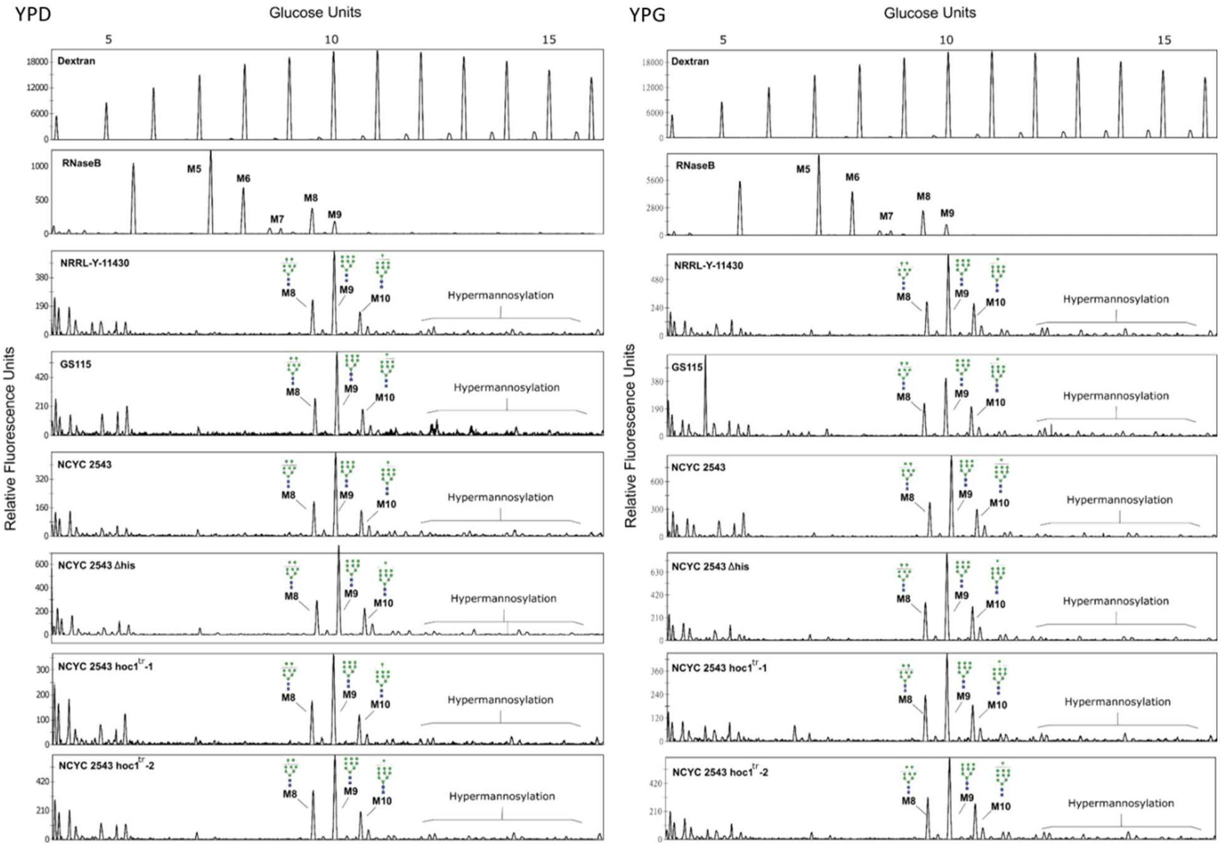


**Supplementary Figure 2. DSA-FACE profiles of the cell wall mannoproteins of the different strains grown on YPD or YPG.** N-glycan profiles of the cell wall mannoproteins when the strains were grown on YPD (left panels) vs. on YPG (right panels). The predominant peaks are $Man_8GlcNAc_2$ (M8), $Man_9GlcNAc_2$ (M9), and $Man_{10}GlcNAc_2$ (M10).

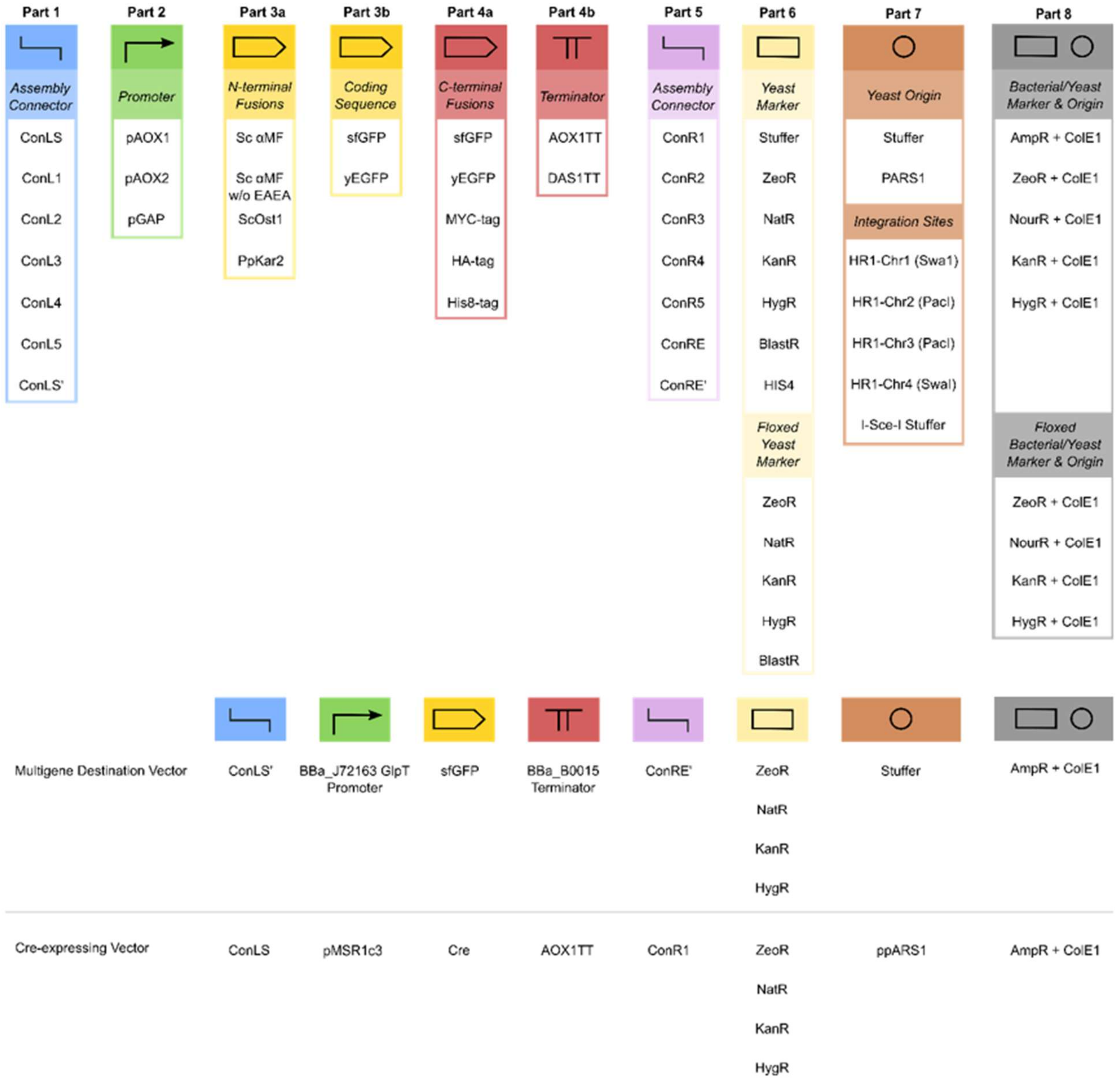


**Supplementary Figure 3. Overview of the available elements for the different Parts of the MoClo toolbox.** For each Part, the available elements are depicted. To generate multigene destination vectors or co-expressing vectors, additional parts are available. All plasmids and plasmid maps are available at the Belgian Co-ordinated Collections of Micro-organisms (BCCM)/GeneCorner Plasmid Collection (http://bccm.belspo.be/about-us/bccm-genecorner).

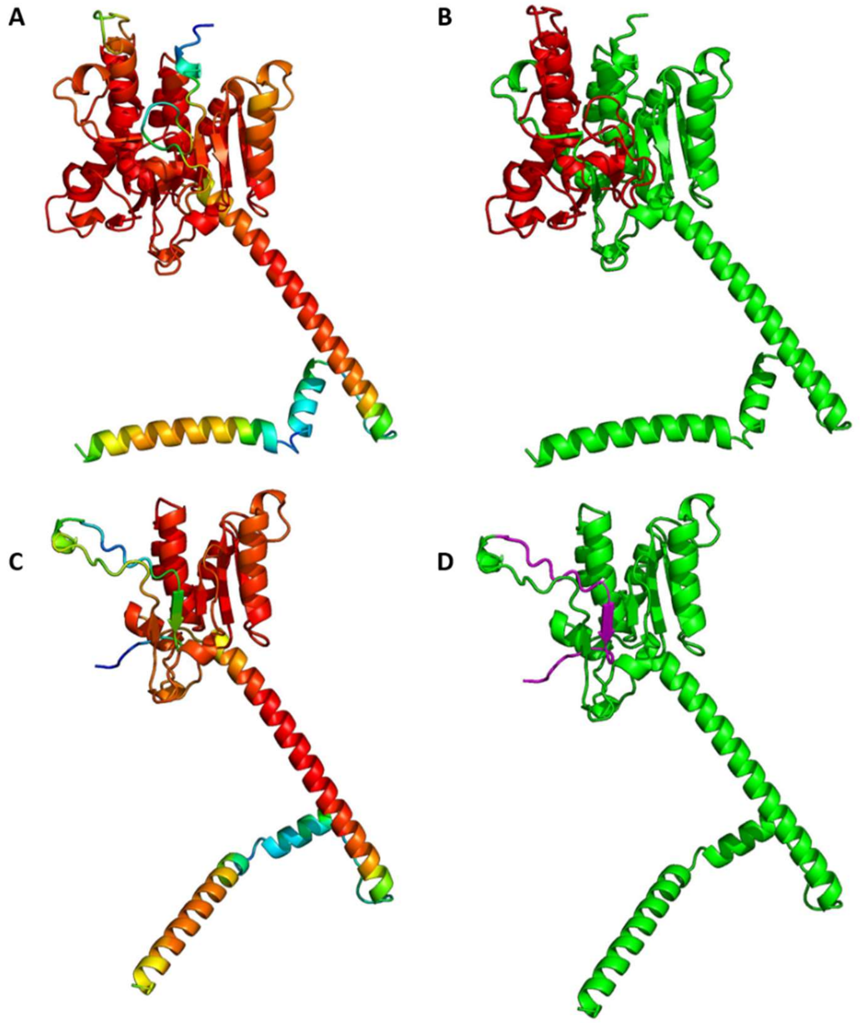


**Supplementary Figure 4. AlphaFold 2 models of the type strain Hoc1p (panels A and B) and of the NRRL Y-11430 lineage derived strain (panels C and D).** The structures in the left panels (A and C) are coloured according to AlphaFold's pLDDT output (prediction reliability score from 0-blue to 100-red). In the right panels (B and D), the wild type C-term piece of sequence (in red) that is replaced by the truncated alternative C-term (magenta) in the NRRL Y-11430 Hoc1p are depicted. Note AlphaFold's prediction that the alternative C-term can still complete the core beta-sheet of the Hoc1p fold (necessarily with low confidence given the absence of multiple sequence alignment data for this alternative C-term).
